# eIF3 interactions with miRNAs involved in translational control of early T cell activation

**DOI:** 10.64898/2026.06.17.732961

**Authors:** Pooja Mukherjee, Jiwon Shin, Cynthia Hermosillo, Chi Yin Ching, Yumi Koga, Aisha Haley Bianchi, Dasmanthie De Silva, Jamie H. D. Cate

## Abstract

Early T cell activation induces extensive remodeling of the cellular transcriptome and proteome. We previously showed using a transcriptome-wide crosslinking approach that human translation initiation factor eIF3 directly interacts with a select set of immune-related mRNAs shortly after T cell activation. We also found that eIF3 binding to the 3′-untranslated regions (3′-UTRs) of the *TCRA* and *TCRB* mRNAs encoding the T cell receptor alpha and beta subunits dynamically regulates a burst in their translation. MicroRNAs (miRNAs) add an additional layer of regulation by fine-tuning both mRNA and protein expression. Although miRNA expression is dynamically regulated during T cell activation, how miRNAs interact with core regulatory pathways in primary T cells remains poorly understood. Here, we reexamined the eIF3-RNA crosslinking experiments in Jurkat cells and probed eIF3 function to investigate miRNA-mediated translational regulation during T cell activation. We found that eIF3 interacts with multiple mature miRNAs in activated Jurkat cells, including members of the miR-17∼92 cluster. These interactions also occur in primary T cells, as shown by RNA immunoprecipitation followed by qPCR (RIP-qPCR). Knocking out the miR-17∼92 cluster led to a delay in ILR2A (CD25) cell surface expression in Jurkat cells and reduced activation-associated cell size increase in primary T cells during early T cell activation. In parallel, we used mass spectrometry to identify eIF3-interacting factors in activated primary T cells, and surprisingly found no evidence of Argonaute binding. Together, these findings provide evidence that eIF3-miRNA interactions may play an unappreciated role in translational control during early T cell activation.

## Introduction

T cell activation is accompanied by a rapid and sustained increase in both transcription and translation that supports the activated state over extended periods (Mukherjee et al., 2025; Wolf et al., 2020). This process includes induction of cell surface markers, cytokine secretion, and proliferation, and unfolds across distinct temporal phases (Cibrián and Sánchez-Madrid, 2017; Peng et al., 2023; Weerakoon et al., 2024). Translational output increases rapidly during early activation and remains elevated during proliferative phases, priming activated T cells for rapid division. Notably, the relationship between mRNA abundance and protein expression is highly dynamic during this process (Mukherjee et al., 2025; Wolf et al., 2020). These observations underscore the central role of translational regulation in shaping the activated T cell proteome and highlight post-transcriptional control as a key determinant of functional outcomes during immune activation. Consistent with this view, previous work from our laboratory demonstrated that human translation initiation factor eIF3 directly interacts with a select set of immune-related mRNAs, including transcripts encoding the T cell receptor alpha and beta subunits (TCRA and TCRB, respectively), and regulates a burst of their expression at the level of translation (De Silva et al., 2021). More broadly, we and others have shown that direct eIF3-mRNA interactions contribute to translational regulation across multiple cellular contexts, supporting a model in which eIF3 functions as an active regulatory hub rather than a passive component of the core initiation machinery (Lamper et al., 2020; Lee et al., 2015; Volta et al., 2021).

MicroRNAs (miRNAs) are another key post-transcriptional regulator in T cells that fine-tune protein expression programs during activation. Although global miRNA activity decreases at later stages of activation (Bronevetsky et al., 2013), specific miRNAs are induced in both CD4⁺ and CD8⁺ T cells, and subsets of miRNAs display dynamic, stage-specific expression during early activation (Mukherjee et al., 2025; Rodríguez-Galán et al., 2018; Xu et al., 2023). Among these, the miR-17∼92 cluster has emerged as a central regulator of immune function (Dölz et al., 2022; Jiang et al., 2011). This highly conserved cluster is encoded within the non-coding host gene *MIR17HG* and produces six mature miRNAs from a single polycistronic transcript (Mendell, 2008). Germline deletion of the miR-17∼92 family results in embryonic lethality, reflecting essential roles in the development of multiple tissues, including the immune system (Ventura et al., 2008). Cell-type-specific perturbations further demonstrate that loss of miR-17∼92 in B cells impairs antibody responses, whereas B-cell-specific overexpression drives lymphomagenesis (Jin et al., 2013a; Sandhu et al., 2013). Mechanistically, miR-17∼92 primarily regulates gene expression at the level of translation, at least in B cells, providing insight into miRNA functional specificity (Jin et al., 2017). In immune contexts, miR-17∼92 promotes differentiation of pathogenic T cell subsets and influences B cell populations, contributing to autoantibody production, while inhibition of miR-17 ameliorates lupus-like chronic graft-versus-host disease (Wu et al., 2018). In T cells, miR-17 and miR-19b are key regulators of Th1 responses (Jiang et al., 2011), and induction of miR-17∼92 downstream of T cell receptor (TCR) and CD28 signaling can compensate for CD28 deficiency, positioning the cluster as a central mediator of T cell activation (Dölz et al., 2022).

Canonical models of microRNA (miRNA) function describe miRNAs acting within Argonaute-containing miRNA-induced silencing complexes (miRISCs) to repress gene expression through sequence-specific interactions that promote translational inhibition and/or mRNA decay (Bartel, 2018; Jonas and Izaurralde, 2015). In animal cells, this repression most commonly arises from partial complementarity between the miRNA 5′ seed region and target transcripts, leading to mRNA deadenylation and decapping. In contrast, near-perfect miRNA-target pairing can trigger Argonaute-mediated endonucleolytic cleavage. Extensive pairing beyond the seed region that is unable to support slicing can instead promote miRNA destabilization through target-directed miRNA degradation (TDMD) (Buhagiar and Kleaveland, 2024; Iwakawa and Tomari, 2022). Thus, the extent and architecture of miRNA-target pairing play a central role in determining regulatory outcomes.

While this framework has provided foundational insight into miRNA-mediated regulation, it does not fully account for the diversity, context dependence, and regulatory flexibility observed across cell types and physiological states (Jens and Rajewsky, 2015; Stavast and Erkeland, 2019). Accumulating evidence indicates that miRNA activity can be modulated by RNA-binding proteins and higher-order ribonucleoprotein assemblies that influence RNA stability, localization, and translational competence (Kedde et al., 2007; Srikantan et al., 2012). Beyond canonical repression, miRNAs have been shown to exert non-traditional regulatory effects, including positive regulation of gene expression (Stavast and Erkeland, 2019; Valinezhad Orang et al., 2014). In some contexts, miRNAs localize to the nucleus and activate transcription through promoter interactions (Valinezhad Orang et al., 2014), while in others they enhance translation by targeting non-seed elements such as AU-rich regions (Vasudevan et al., 2007) or 5′ terminal oligopyrimidine (TOP) motifs within untranslated regions (Ørom et al., 2008). miRNAs can also act as decoys, sequestering ribonucleoproteins in a seed-independent manner and indirectly promoting translation of target mRNAs (Eiring et al., 2010). Together, these findings support an expanded view of miRNA function that extends beyond Argonaute-centric repression and raise the question of whether miRNAs could engage components of the translation initiation machinery to achieve context-specific regulation, particularly during dynamic processes such as T cell activation.

In this study, we examined how miRNA-mediated regulation intersects with translational control during early T cell activation. We first reexamined eIF3 crosslinking to RNAs early after Jurkat cell activation (De Silva et al., 2021), and found that eIF3 crosslinks extensively to mature miRNAs, including members of the miR-17∼92 cluster. We then used affinity purification of eIF3-RNA complexes from primary T cells to validate the crosslinking experiments. We also used mass spectrometry of eIF3-associated factors in activated primary T cells, which suggest that eIF3-miRNA interactions occur independently of Argonaute proteins. Functional perturbation of miRNAs in the miR-17∼92 cluster affects T cell activation, consistent with a role for miRNA-eIF3 interactions in shaping activation-associated translational responses. Together, this work provides evidence that eIF3 engagement of miRNAs may represent an additional layer of post-transcriptional regulation during immune cell activation.

## Results

### eIF3 interactions with miRNAs in Jurkat and primary T cells

We previously identified RNAs bound to eIF3 in Jurkat cells activated with Phorbol 12-myristate 13-acetate (PMA) and ionomycin using Photoactivatable Ribonucleoside-Enhanced Crosslinking and Immunoprecipitation (PAR-CLIP) and 4-thiouridine (De Silva et al., 2021; Hafner et al., 2010). PAR-CLIP experiments with 4-thiouridine incorporated into RNAs provide a diagnostic signature of T to C transitions in the resulting DNA mapped reads (De Silva et al., 2021; Hafner et al., 2010). Using this method, we identified multiple immune-related mRNAs important for robust T cell activation including the *TCRA* and *TCRB* mRNAs encoding the alpha and beta subunits of the T cell receptor. The original PAR-CLIP data analysis used the PARalyzer program as implemented in the PARpipe workflow (De Silva et al., 2021; Mukherjee et al., 2019). The data processed in this way revealed eIF3 crosslinking to miRNAs, but was too conservative in handling PCR duplication artifacts, by relying first on unique molecular identifier (UMI) sequences, and then the default PARalyzer assumption of identical reads being duplicates. For miRNAs which have narrow size distributions and few U residues available for crosslinking (and therefore T to C transitions), this likely overly compressed eIF3 crosslinking reads to miRNAs.

We therefore reprocessed the eIF3 PAR-CLIP data to rely solely on the UMI sequences incorporated during library preparation (Methods). When processed using the UMI sequences, we observed multiple miRNAs extensively crosslinked to eIF3, including nearly all of the miRNAs encoded in the *MIR17HG* cassette (Figure 1A, Figure 1–figure supplement 1). Notably, eIF3 crosslinking is much higher to the guide strand of each miRNA, as indicated by miRBase (Kozomara and Griffiths-Jones, 2011). We compared the eIF3 crosslinking pattern of T to C transitions with previously published Argonaute 2 (AGO2) PAR-CLIP data (Hafner et al., 2010) and observed a strikingly similar but not identical pattern for eIF3 subunits EIF3A and EIF3B and AGO2 (Fig. 1B; Figure 1–figure supplement 2, Supplementary file 1). By contrast, subunit EIF3D showed a different pattern of T to C transitions in the corresponding miRNAs (Fig. 1B; Figure 1–figure supplement 2). Next, we aimed to identify predicted mRNA targets of these miRNAs from among the mRNAs that crosslink to eIF3 in activated Jurkat cells in the PAR-CLIP dataset. We observed substantial overlap between predicted targets and eIF3-associated transcripts, with the strongest overlap for miR-19a-3p (Figure 1–figure supplement 3A). Gene Ontology analysis of the overlapping targets revealed enrichment for pathways related to T cell functions, among others (Figure 1–figure supplement 3B).

**Figure 1.**
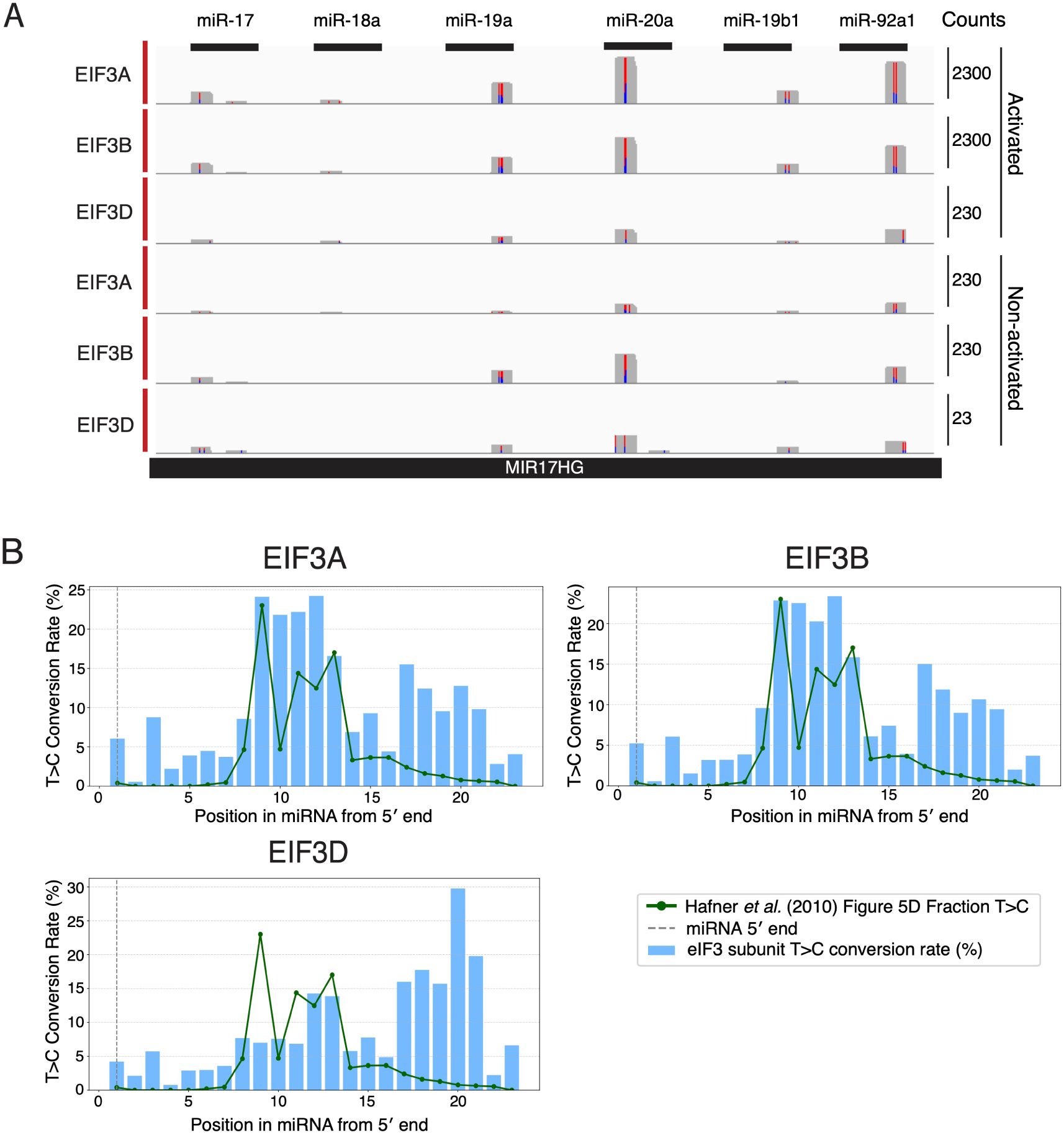
eIF3 crosslinks with specific microRNAs in activated Jurkat cells. (A) Crosslinking of the indicated eIF3 subunits across all mature miRNAs of the miR-17∼92 cassette in activated and unactivated Jurkat cells (replicate 1). (B) T>C conversion profiles from PAR-CLIP data plotted relative to the miRNA 5′ end for EIF3A, EIF3B, and EIF3D in activated Jurkat cells. Blue bars represent the T>C conversion rate (%) at each nucleotide position along the miRNA, the green line represents the published AGO2 T>C conversion profile from Hafner et al (2010) and the grey dotted line represents the miRNA 5′ end.

To examine whether eIF3 also associates with miRNAs in primary T cells, we performed RNA immunoprecipitation (RIP) using an antibody against EIF3B and RT-qPCR to detect bound miRNAs (Figure 2A). We prepared lysates from unactivated T cells and T cells activated for 5 hours (5h) using anti-CD3/anti-CD28 antibodies and affinity purified eIF3 from these samples. We washed the eIF3-bound beads under low-salt conditions to preserve RNA-protein interactions and to maintain the integrity of the multisubunit eIF3 complex. Under these conditions, EIF3B was efficiently immunoprecipitated, indicating similar IP efficiency across activation states (Figure 2B, Figure 2–figure supplement 1). We then isolated RNA from EIF3B immunoprecipitates for qPCR analysis (Methods). Notably, multiple members of the miR-17∼92 cluster were enriched in EIF3B IP samples relative to IgG controls (Figure 2C, Figure 2–figure supplement 2, Supplementary file 2). In contrast to the PAR-CLIP experiment in Jurkat cells, we did not observe a clearly higher enrichment of these miRNAs at 5h compared with unactivated cells.

**Figure 2.**
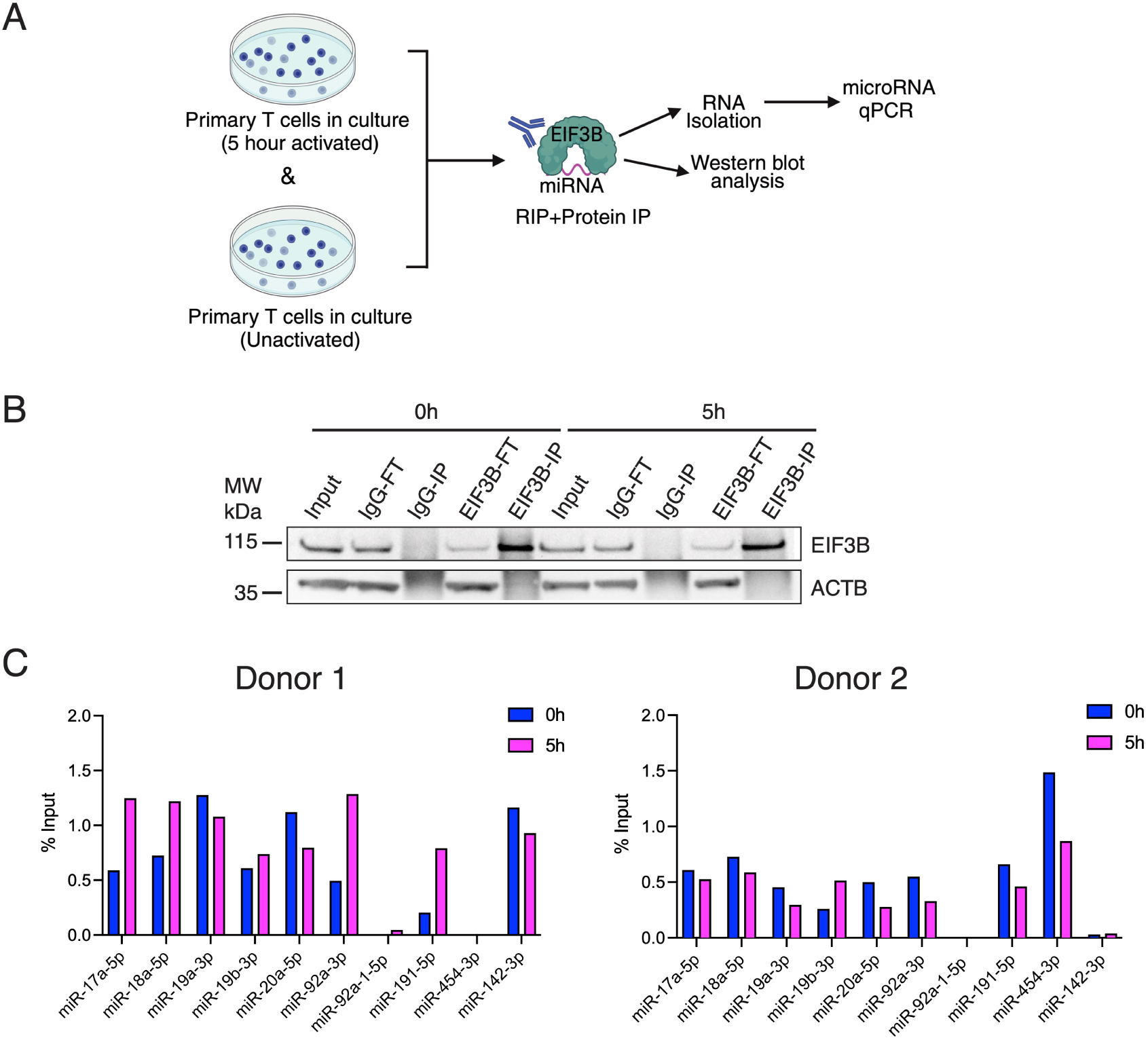
EIF3B interacts with the miR-17∼92 cassette in primary T cells. (A) Schematic of the EIF3B RNA immunoprecipitation (RIP)-miRNA qPCR workflow in unactivated and 5 hour activated primary T cells. (B) Western blot analysis of EIF3B immunoprecipitation from unactivated and 5 hour activated primary T cells. Input, flowthrough (FT), and immunoprecipitated (IP) fractions are shown, with ACTB as a loading control. (C) RIP-qPCR analysis of miRNAs from the miR-17∼92 cassette enriched in EIF3B immunoprecipitation from unactivated and 5 hour activated primary T cells (Donor 1 and Donor 2), plotted as % Input (% Input EIF3B minus % Input IgG).

### Role of the miR-17∼92 cluster in T cell activation

Due to the observed binding of miR-17∼92 cluster miRNAs to eIF3, we investigated the role of the miR-17∼92 cluster in T cell activation. We first generated Jurkat cell lines carrying a complete deletion of the miR-17∼92 genomic locus (Figure 3A). Single-cell clones were isolated and screened using flanking PCR primers, and two independent homozygous knockout clones were selected for downstream analyses. These clones were used for all phenotypic and transcriptional experiments using Jurkat cells (Figure 3–figure supplement 1A-1B, Supplementary file 3). T cell activation proceeds through early and late phases (Marchingo and Cantrell, 2022; Weerakoon et al., 2024), marked by distinct surface expression patterns, with CD69 induced rapidly after stimulation (Cibrián and Sánchez-Madrid, 2017; Gorabi et al., 2020) and CD25 (IL2RA) upregulated at later stages as part of the IL-2 receptor (Peng et al., 2023). After anti-CD3/anti-CD28 stimulation of the Jurkat cells, flow cytometry revealed that deletion of miR-17∼92 resulted in a reproducible reduction in CD25 expression in both knockout clones, with the strongest effect observed at 12h post-activation. In contrast, effects on CD69 were more modest (Figure 3B, Figure 3–figure supplement 1C). Re-expression of the miR-17∼92 cluster in knockout cells restored CD25 surface expression, confirming that the observed activation defect was specifically attributable to miR-17∼92 loss (Figure 3–figure supplement 2).

**Figure 3.**
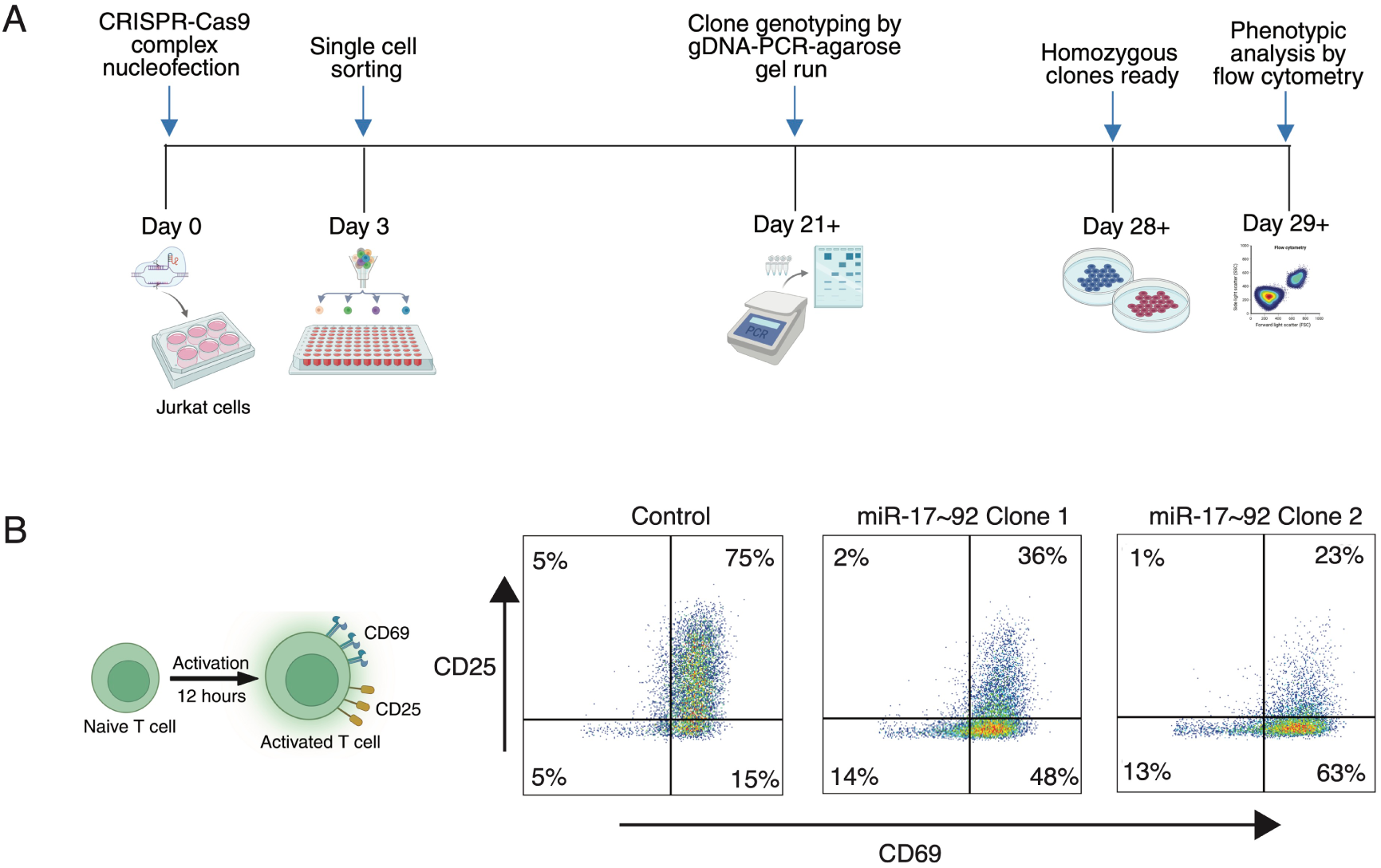
Generation and phenotypic analysis of miR-17∼92 knockout Jurkat cells. (A) Schematic of the CRISPR-Cas9 workflow used to generate miR-17∼92 knockout Jurkat cell clones. (B) Flow cytometry analysis of CD25 and CD69 expression following activation of unedited Jurkat cells and miR-17∼92 knockout clones for 12 hours. Representative plots are shown, with quadrant percentages indicated.

To define early transcriptional changes associated with miR-17∼92 deletion, we performed mRNA sequencing on wild-type and knockout clones at 0h (unactivated) and 5h following activation using anti-CD3/anti-CD28 antibodies. Principal component analysis (PCA) showed clustering of biological replicates and variation associated with genotype and activation state (Figure 4A). Differential expression analysis revealed substantial differences in transcript abundance between wild-type and knockout cells at both time points. Notably, at both 0h and 5h, the number of downregulated genes exceeded the number of upregulated genes in the knockout clones (Figure 4B, Figure 4–figure supplement 1A, Supplementary file 4), despite the canonical role of miRNAs as negative regulators of gene expression. This suggests there may be indirect or context-dependent effects of miR-17∼92 loss in Jurkat cells. To identify reproducible activation-associated changes, we focused on differentially expressed genes shared between the two knockout clones at 5h post-activation (Figure 4C, Figure 4–figure supplement 1B, Supplementary file 4). Gene ontology analysis of this overlapping gene set revealed enrichment for multiple T cell-related pathways, including T cell activation and cytokine signaling (Figure 4D, Supplementary file 4). Consistent with these findings, analysis of individual T cell activation genes within this set revealed altered expression of transcripts associated with CD25 and CD44 regulation, among others (Figure 4E, Supplementary file 4), correlating with the reduced surface expression of CD25 observed in miR-17∼92 knockout cells and recapitulating previously reported roles of the miR-17∼92 cluster in T cell activation (Dölz et al., 2022).

**Figure 4.**
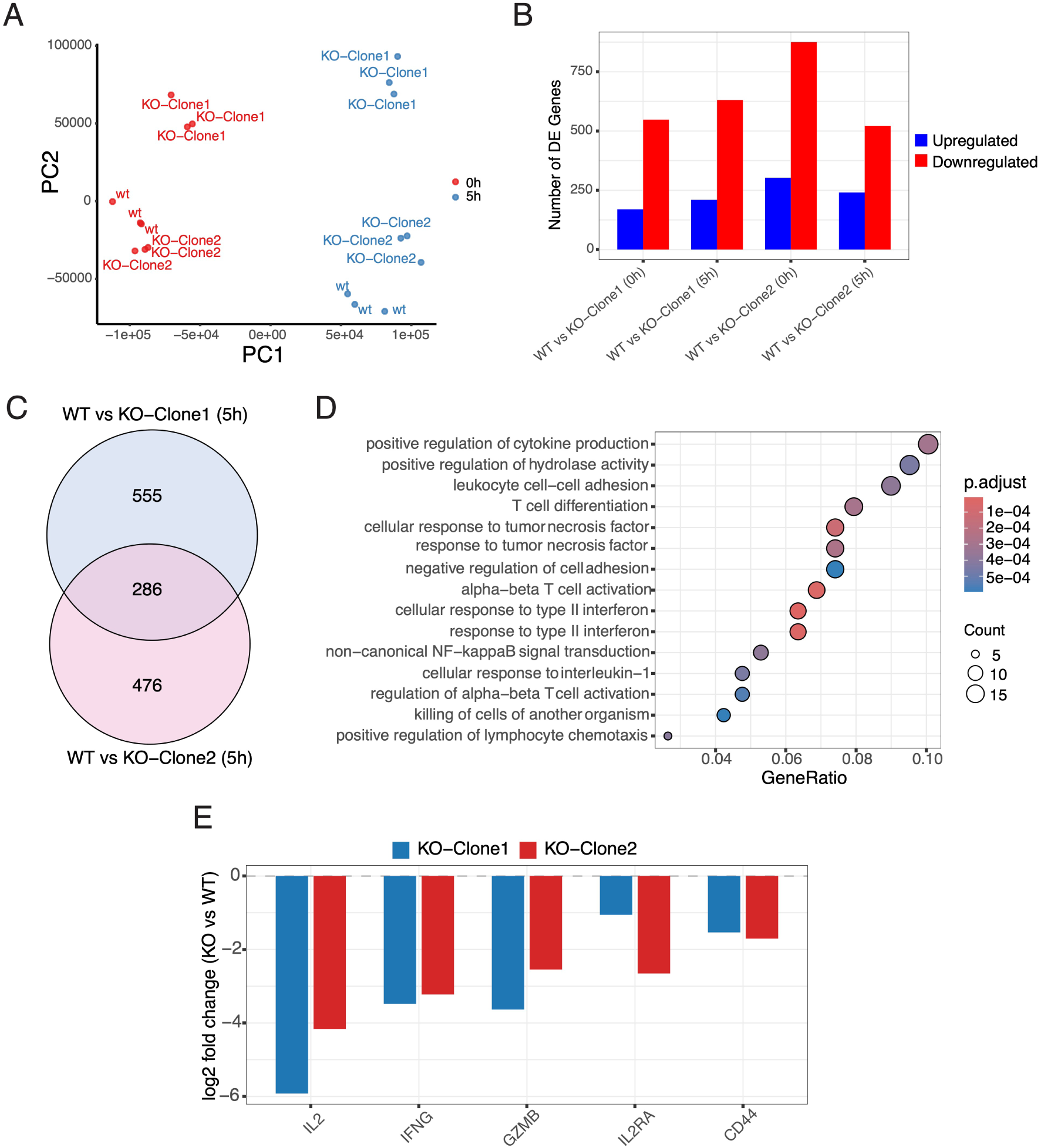
Transcriptomic analysis of miR-17∼92 knockout Jurkat cell clones. (A) Principal component analysis (PCA) of RNA-seq samples from Wild-type (WT) and miR-17∼92 knockout Jurkat clones under unactivated and 5 hour activation conditions. (B) Number of differentially expressed (DE) genes identified in WT versus miR-17∼92 knockout clones under unactivated and 5 hour activation conditions (|log2 fold change| > 1). (C) Overlap of DE genes between miR-17∼92 knockout clones at 5 hour activation. (D) Gene Ontology (GO) analysis of DE genes at 5 hour activation. (E) Expression changes of selected T cell-related genes at 5 hour activation, shown as log2 fold change relative to WT.

To extend these findings to nontransformed cells, we disrupted the miR-17∼92 cluster in primary human T cells using CRISPR-Cas9 (Figure 5A-B). Two independent pairs of flanking guide RNAs were tested in four different combinations to excise the miR-17∼92 locus. Editing efficiency was assessed by flanking PCR and agarose gel analysis, and the guide combination yielding the highest deletion efficiency was selected for downstream experiments (Figure 5B, Supplementary file 3). Sequencing of the PCR products revealed approximately 77-81% editing efficiency in bulk T cell populations (Figure 5–figure supplement 1A). Loss of miR-17∼92 was further confirmed by miRNA-specific qPCR, which showed a marked reduction in mature miR-17∼92 cluster members compared with wild-type cells (Figure 5C, Supplementary file 5). Because primary T cells are not easily clonally expanded following genome editing, all subsequent analyses were performed on bulk-edited populations. Upon activation using anti-CD3/anti-CD28 antibodies, miR-17∼92-edited T cells displayed a reduction in cell size relative to wild-type cells, as assessed by forward scatter measurements (Figure 5D). This phenotype was notable given that T cell activation is normally accompanied by an increase in cell size associated with blast formation (Waugh et al., 2023). In contrast to the effects observed in Jurkat knockout clones, no consistent reduction in CD25 surface expression was detected in bulk-edited primary T cells (Figure 5–figure supplement 1B). These results suggest that deletion of miR-17∼92 in heterogeneous primary T cell populations is sufficient to impair activation-associated cell growth, while effects on CD25 expression may be more specific to Jurkat cells.

**Figure 5.**
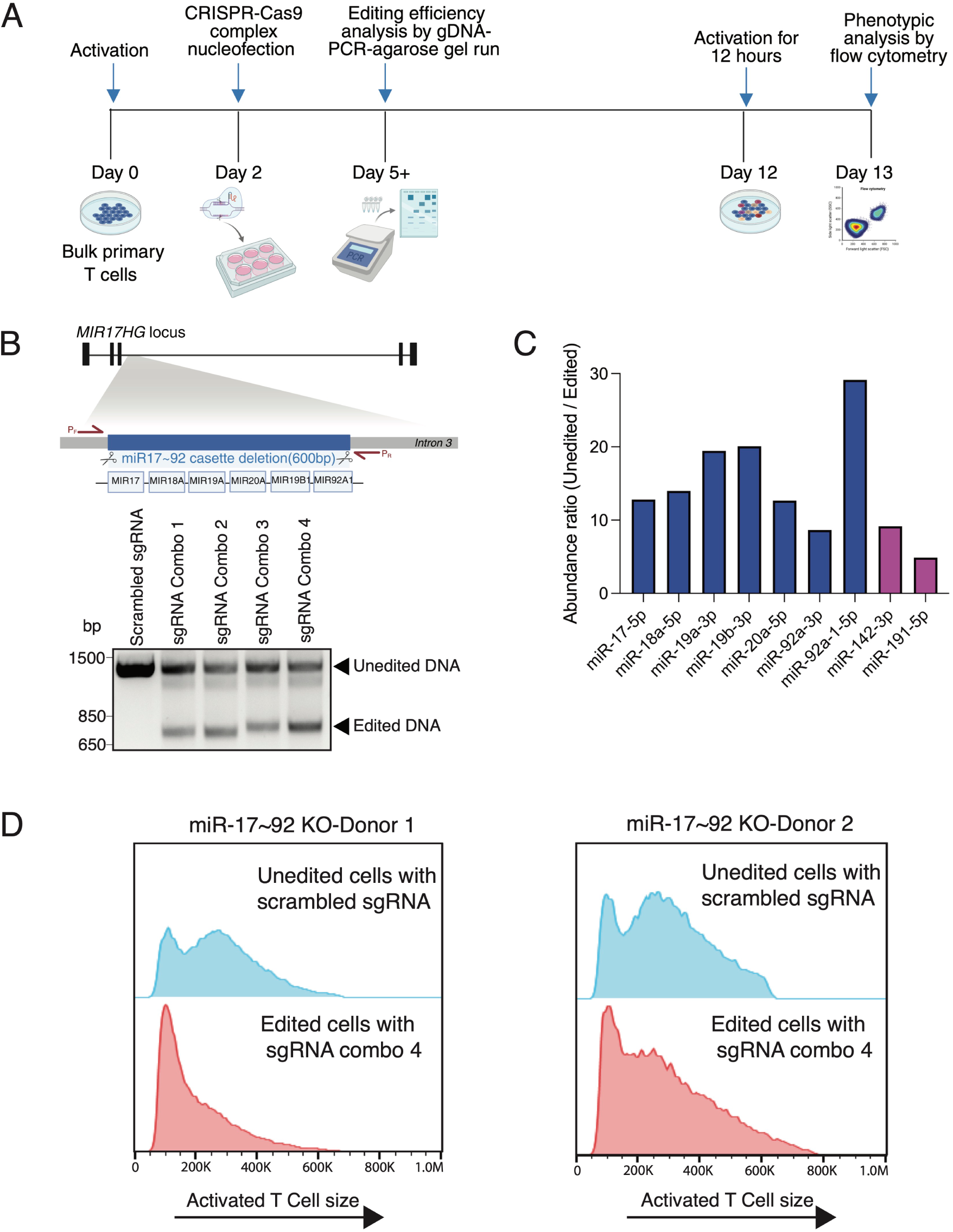
CRISPR-Cas9 editing and phenotypic analysis of miR-17∼92 in primary T cells. (A) Schematic of the CRISPR-Cas9 workflow used to edit the miR-17∼92 locus in bulk primary T cells. (B) Schematic of the miR-17∼92 locus and PCR-based genotyping strategy, with representative agarose gel showing unedited (scrambled sgRNA) and edited DNA (sgRNA combination 1-4) products for the indicated sgRNA combinations. (C) qPCR analysis comparing miRNA expression in unedited and edited bulk primary T cells (sgRNA combination 4), shown as abundance ratio of edited and unedited cells. (D) Flow cytometry analysis of activated T cell size, measured by forward scatter area (FSC-A), in unedited and edited live primary T cells from two donors (Donor 1 and Donor 2).

### eIF3-protein interactions in activated T cells

We wondered whether components of the RNA-induced silencing complex or other proteins related to miRNA function might associate with eIF3 in T cells. In order to identify proteins associated with eIF3, we performed eIF3 immunoprecipitation using the antibody against EIF3B as above, with or without RNase treatment, followed by mass spectrometry (IP-MS) (Figure 6A). Under all conditions tested, EIF3B was efficiently immunoprecipitated, and multiple core subunits of the eIF3 complex were consistently recovered across replicates (Figure 6B, Supplementary file 6), confirming robust and reproducible isolation of the intact eIF3 complex. Interestingly, AGO2 was not detected in the EIF3B IP-MS datasets and by western blot (Figure 6C, Figure 6–figure supplement 1A). This was true for both RNase treated and untreated samples, in both unactivated and T cells activated by anti-CD3/anti-CD28 antibodies for 5h. Consistent with this observation, gene ontology analysis did not reveal enrichment for canonical RISC components or core miRNA pathway proteins (Figure 6–figure supplement 1B, Supplementary file 6). Together, these results indicate that EIF3B does not detectably associate with AGO2 under the conditions tested, although it does engage a distinct set of RNA-associated factors through RNA-dependent interactions that are separable from canonical RISC complexes (Figure 6D, Supplementary file 6).

**Figure 6.**
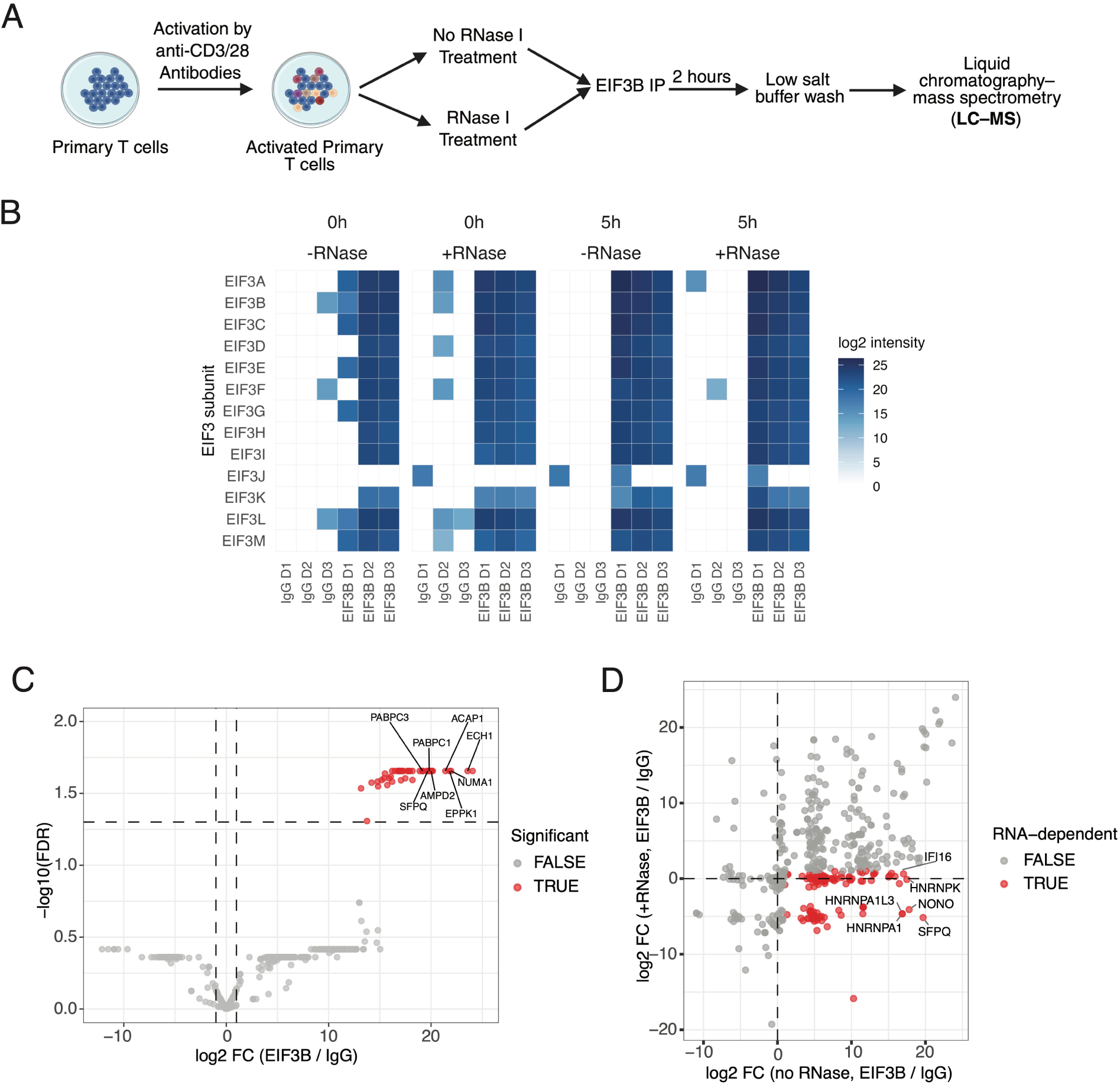
Proteomic analysis of EIF3B-associated complexes in activated primary T cells. (A) Schematic of the experimental workflow for EIF3B immunoprecipitation followed by mass spectrometry analysis in unactivated and 5 hour activated primary T cells, with or without RNase I treatment. (B) Heatmap showing relative abundance of eIF3 complex subunits detected across conditions at unactivated and 5 hour activation, with and without RNase treatment. (C) Volcano plot of proteins enriched in EIF3B immunoprecipitation compared to IgG control at 5 hour activation in the absence of RNase treatment (eIF3 complex subunits removed). (D) Comparison of protein enrichment in EIF3B immunoprecipitation with and without RNase treatment at 5 hour activation, highlighting RNase-sensitive interactions.

## Discussion

Previously we showed eIF3 regulates the translation of mRNAs central to early T cell activation, including those encoding T cell receptor subunits TCRA and TCRB (De Silva et al., 2021). Here, we found that eIF3 also binds miRNAs in T cells, including those in the miR-17∼92 cluster (Figures 1-2). Specifically, we observed preferential crosslinking to the dominant or guide strands across all miRNAs within the miR-17∼92 cluster. Notably, the PAR-CLIP data also reveal an eIF3 subunit-specific architecture within the eIF3-miRNA interface. The T to C transition patterns for EIF3A and EIF3B are strikingly similar to those previously observed for AGO2 (Hafner et al., 2010), suggesting these eIF3 subunits may also bind miRNA-mRNA complexes engaged through miRNA 5′ seed pairing. Such miRNA-mRNA interactions would be consistent with eIF3’s known binding to structured RNA elements (Kieft et al., 2001; Lamper et al., 2020; Lee et al., 2015). In contrast, EIF3D exhibits a distinct crosslinking profile, supporting the model that eIF3 subunits can exert specialized, non-canonical functions in translational control beyond the core initiation scaffold. EIF3D extends from the eIF3 core scaffold (Brito Querido et al., 2020; des Georges et al., 2015; Simonetti et al., 2020) and is therefore well positioned to mediate interactions beyond the central eIF3 complex. Beyond its role in general translation initiation, EIF3D has been shown to carry out non-canonical functions in translational control, including selective mRNA recognition and cap-binding activity that can bypass EIF4E-dependent initiation (Lee et al., 2016; Volta et al., 2021), suggesting a role in transcript-specific regulatory programs.

While eIF3-miRNA binding increased upon Jurkat cell activation, qPCR analysis in primary T cells demonstrated enrichment in both unactivated and activated states. Furthermore, the relative enrichment of mature miR-17∼92 cluster miRNAs observed by qPCR was more modest than the PAR-CLIP signal. Several methodological factors could account for this discrepancy. First, the qPCR approach in primary cells relied on non-crosslinked immunoprecipitation, which is inherently more qualitative and prone to the dissociation of transient interactions compared to the covalent stabilization provided by PAR-CLIP. Second, variations in primer efficiency across different miRNA targets may have hindered precise comparative quantification. Finally, the use of low-salt wash conditions, while necessary to preserve transient miRNA-eIF3 interactions, may have increased the recovery of non-specific small RNAs, thereby narrowing the observed enrichment margin. Despite these quantitative challenges, the reproducible detection of miR-17∼92 enrichment across independent experiments and platforms supports a specific biochemical association. Together, these observations suggest that miRNA association with eIF3 reflects regulatory roles that extend beyond canonical models in which miRNAs function exclusively within Argonaute-centered effector complexes (Jens and Rajewsky, 2015; Stavast and Erkeland, 2019). Future experiments in primary T cells will be required to identify the possible roles for eIF3-miRNA interactions.

A striking enrichment of eIF3 binding to miRNAs encoded in the miR-17∼92 cluster raises the possibility that eIF3 may be responsible for some of the roles of these miRNAs in T cell activation. The miR-17∼92 cluster has previously been shown to play a role in immune regulation across multiple contexts. For example, they play roles in antibody responses and lymphomagenesis in B cells (Jin et al., 2013b; Sandhu et al., 2013). In T cells, individual members such as miR-17 and miR-19b regulate Th1 responses (Jiang et al., 2011), and induction of the cluster downstream of TCR and CD28 signaling can compensate for CD28 deficiency, underscoring its central role in T cell activation (Dölz et al., 2022). Here, we discovered additional roles for these miRNAs in early T cell activation. In Jurkat cells, deletion of *MIR17HG* led to a decrease in CD25 expression 12 hours after activation. In primary T cells, deletion of *MIR17HG* caused a decrease in cell size after activation, consistent with a blunted activation response. Future experiments will be required to test whether eIF3 plays a direct role in the miR-17∼92 cluster mediated phenotypes we observed here (Figure 7).

**Figure 7.**
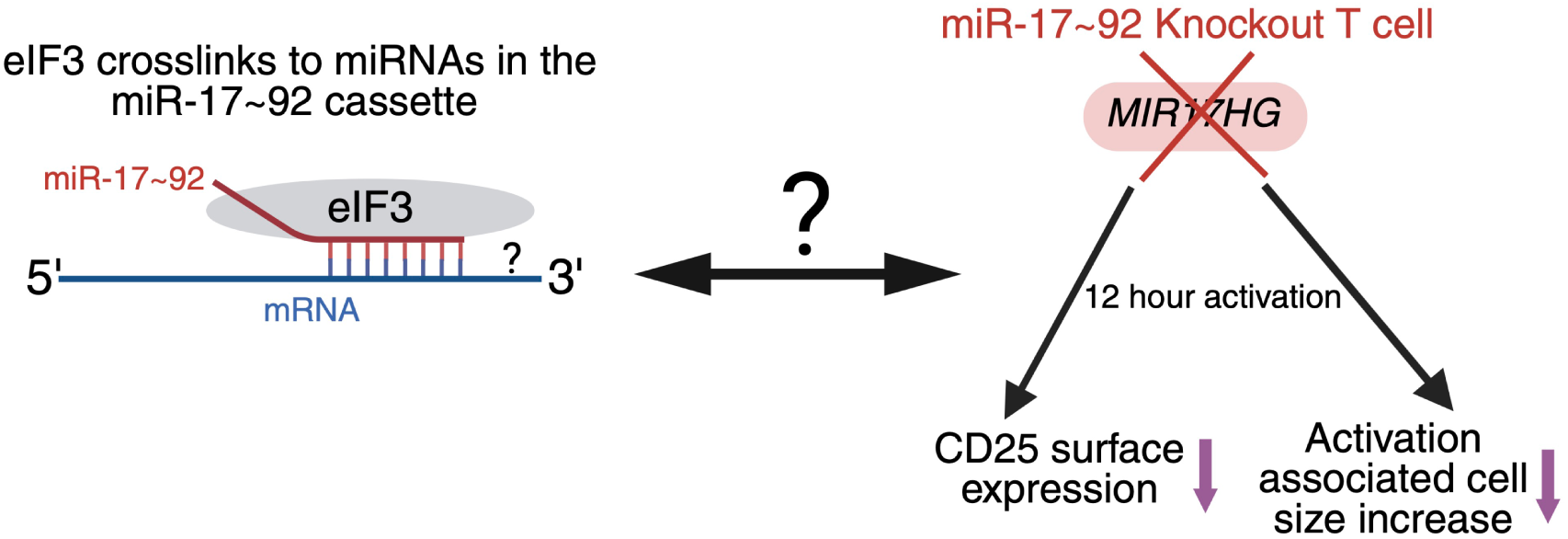
A Model for eIF3 and miR-17∼92-Dependent Control of T Cell Activation Phenotypes. Schematic summarizing observed molecular interactions and T cell knockout phenotypes. (Left) Data show binding of EIF3 to the mature miRNAs of the miR-17∼92 cassette. (Right) Genetic deletion of the *MIR17HG* gene in activated T cells leads to two distinct cellular defects: reduced CD25 surface expression and reduced activation-associated cell size after 12 hours of activation. The double-headed arrow marked with a question mark highlights the mechanistic question of how the direct eIF3-miRNA interaction functionally regulates these specific T cell activation outcomes. The question mark on the mRNA indicates that the presence and identity of an mRNA target in the eIF3-miRNA complex have not been established.

The action of miRNAs in regulating mRNA translation and miRNA turnover is regulated by AGO2 and the RISC complex. We tested whether eIF3 association with miRNAs might involve interactions with AGO2. However, the crosslinking pattern of eIF3 to miRNAs reflected in the T to C transitions would seemingly preclude other protein-miRNA interactions for steric reasons, given the extensive miRNA-protein interactions seen in AGO2 complexes (Schirle and MacRae, 2012). Indeed, we were unable to detect AGO2 or any RISC component in our EIF3B IP-MS analyses. This suggests that eIF3 binding to miRNAs occurs independently of the canonical miRNA silencing pathways, and raises the question of whether other trans-acting factors might contribute to eIF3-miRNA function. One cytoplasmic protein enriched to the same extent as other eIF3 subunits in the PI-MS analyses is ACAP1. ACAP1 is a GTPase-activating protein (GAP) for ADP ribosylation factor 6 (ARF6), which is involved in clathrin-dependent export of proteins from recycling endosomes to the cell surface. Notably, increased transient expression of the T cell receptor mediated by eIF3 early after T cell activation is dependent on membrane-localization of eIF3 (De Silva et al., 2021), suggesting that eIF3 may be enriched near recycling endosomes in activated T cells. Notably, analysis of RNase-treated and RNase-untreated samples revealed the presence of multiple nuclear-localized proteins. Because total cell lysates were used in these experiments, we cannot rule out nonspecific binding of these proteins.

Given evidence for AGO2-independent association of EIF3B with miRNAs, several models for eIF3-miRNA interactions can be proposed. One possibility is a “handoff model”, in which miRNAs transition between distinct protein assemblies as activation progresses. In such a model, miRNAs may initially function within canonical Argonaute-containing complexes and subsequently associate with eIF3 regulatory assemblies under specific cellular conditions. While our study does not directly demonstrate such transitions, some features of T cell activation are consistent with this possibility. Notably, Ago2 has been reported to undergo degradation during T cell activation, suggesting that canonical miRISC activity may be dynamically regulated (Bronevetsky et al., 2013). However, a conceptual challenge for a handoff-based model is how miRNAs could be released from Argonaute proteins without being degraded, as occurs in target-mediated miRNA decay (TDMD) (Buhagiar and Kleaveland, 2024; Farnung et al., 2026; Hiers et al., 2024), in which extensive base pairing between a miRNA and its target leads to miRNA destabilization rather than target repression. Alternatively, eIF3 may selectively associate with a specific subset of miRNAs immediately following their biogenesis and processing, independent of the canonical RNA-induced silencing complex. To test these possibilities, time-resolved parallel CLIP of AGO2 and EIF3B could be employed to track whether specific miRNAs exhibit reciprocal occupancy patterns across these complexes during T cell stimulation. Distinguishing between these models will also require high-resolution mapping of the eIF3-miRNA-target interactome to identify the precise mRNA targets co-captured by eIF3 and miRNAs. These experiments will be essential to determine if this association facilitates a regulatory “switch” from canonical silencing to specialized translational control during T cell activation.

Downstream of these associations, eIF3 could influence miRNA regulatory outcomes in multiple ways. For example, association with eIF3 may support miRNA-mediated repression of specific targets, facilitate context-dependent activation, or act as a buffering mechanism that limits miRNA access to certain targets, thereby relieving repression. While our data do not distinguish between these possibilities, they raise the idea that eIF3-associated miRNAs may participate in regulatory modes that differ from canonical Argonaute-centered repression. The relevance of these regulatory possibilities is underscored by the central role of the miR-17∼92 cluster in T cell biology.

Several limitations of this study should be noted. Our analyses primarily establish molecular associations between eIF3 and miRNAs rather than direct mechanistic relationships with mRNA regulation, and the determinants governing miRNA association with eIF3-containing complexes remain undefined. In addition, while we examined both primary T cells and Jurkat cells, further work will be needed to assess the generality of these observations across immune cell types and activation states. Future studies aimed at defining the temporal dynamics, molecular interfaces, and functional consequences of miRNA-eIF3 interactions will be essential. In summary, this work provides evidence that miRNAs implicated in immune regulation associate with eIF3 during early T cell activation. By identifying interactions between eIF3 and mature miRNAs, including members of the miR-17∼92 cluster, our findings suggest that miRNAs engage eIF3-containing regulatory assemblies during periods of rapid cellular remodeling (Figure 7). Our findings expand current models of post-transcriptional regulation in immune cells and highlight new directions for investigating how noncoding RNAs contribute to rapid proteome remodeling during immune activation.

## Methods

### PAR-CLIP data reprocessing

We reprocessed the eIF3 PAR-CLIP data obtained from Jurkat cells (De Silva et al., 2021) to map reads to a more up-to-date human genome annotation, and to improve mapping of reads to microRNAs. Data for sequence reads of the eIF3 PAR-CLIP experiment in activated and nonactivated Jurkat cells were downloaded from the NCBI Sequence Read Archive (SRA) using accession SRP351691. Details of DNA library preparation are given in the NCBI Gene Expression Omnibus, accession GSM5743370. The libraries include 5-nucleotide unique molecular identifiers (UMIs) in the 3′ primer sequence, which were used for PCR deduplication. Briefly, barcoded reads were processed with cutadapt (Martin, 2011) into separate files for samples enriched for EIF3A, EIF3B, or EIF3D crosslinking (De Silva et al., 2021). Each file was then deduplicated using awk and the Unix “sort -u” command, to remove possible PCR amplification artifacts.

After removing the 3′ adapter sequence using cutadapt, reads were mapped to the human genome using Hisat-3n (Zhang et al., 2021). The GRCh38 release 1.04 primary assembly of the human genome was obtained from Ensemble (Cunningham et al., 2022), and used with Hisat-3n, indexed to allow T to C transitions expected in PAR-CLIP reads (hisat-3n-build --base-change T,C). Mapped reads were then filtered using samtools (Danecek et al., 2021) and the Hisat-3n supplied Yf field to remove reads with more than 2 T to C transitions, with the assumption these might be incorrectly mapped reads. The samtools command used was the following:

samtools view -h -e ’[Yf] >= 0 && [Yf] <= 2’

To identify reads that map to miRNAs, we first downloaded the hg38 coordinates for miRNAs from miRBase (Kozomara et al., 2019), filtered for mature miRNAs, and converted these to .bed format. We then used bedtools to identify overlaps between eIF3 PAR-CLIP reads and miRNAs, with at least 90% of the miRNA sequence overlapped, using the command:

bedtools intersect -f 0.9 -u

We then extracted miRNA-intersecting reads from the Hisat-3n and T-to-C transition filtered .bam files, to generate new .bam files containing only miRNA-mapped reads. We used these reads, the mature miRNA coordinates, and the hg38 primary assembly to generate meta-miRNA alignments of all EIF3A, EIF3B, and EIF3D sample reads. Only miRNAs with at least 10 mapped reads in a sample (excluding secondary reads as defined by samtools) were used for the meta-miRNA alignments. These alignments were used to calculate the fraction of T to C transitions at each miRNA position using a custom python script. We then compared the fraction of T to C transitions at each nucleotide position (positions 1 through 23) to those observed for miRNAs crosslinked to Argonaute 2 (Ago2) by Hafner *et al*. (Hafner et al., 2010), in their Figure 5D. For plots of miRNA reads in the Integrated Genomics Viewer (IGV) (Thorvaldsdóttir et al., 2013), reads mapping to multiple locations were randomly assigned genomic positions for the primary read for display in IGV.

### Identification of miR-17∼92 targets crosslinked by eIF3 in Jurkat PAR-CLIP

The eIF3 Jurkat PAR-CLIP cluster dataset was downloaded from NCBI GEO (GSE191306) and analyzed using the file: GSE191306_DeSilva-Jurkat-eIF3-PAR-CLIP-clusters.xlsx. We focused on mature, arm-resolved miRNAs from the miR-17∼92 cluster: hsa-miR-17-5p, hsa-miR-18a-5p, hsa-miR-19a-3p, hsa-miR-20a-5p, hsa-miR-19b-3p, and hsa-miR-92a-3p.

Experimentally validated targets for each miRNA were downloaded from DIANA-TarBase v9, filtering to “Homo sapiens” and restricting experimental support to “Direct” evidence. Targets were exported per miRNA, merged across miRNAs, and deduplicated by gene symbols to generate per-miRNA and union target sets. Predicted targets were obtained from TargetScan 8.0 and filtered to the same six miRNAs (accounting for TargetScan family groupings). Targets were deduplicated by gene symbol, retaining the entry with the most negative Total context++ score, and saved as per-miRNA and union target lists. Overlap between miRNA target genes (validated and predicted) and eIF3-crosslinked genes from the PAR-CLIP dataset was filtered for intersections, producing per-miRNA and union counts of targets detected in the Jurkat eIF3 PAR-CLIP data.

### Primary T cell culture

Peripheral blood mononuclear cells (PBMCs) were isolated from leukopaks (Stemcell) by sequential washing and low-speed centrifugation in MACS/EasySep buffer (PBS supplemented with 2% fetal bovine serum and 1 mM EDTA), with an initial centrifugation at 300 × g for 10 minutes to remove residual plasma, followed by centrifugation at 120 × g for 10 minutes without brake to enrich for PBMCs. Bulk T cells (CD3) are isolated from PBMCs by magnetic selection using EasySep Human T Cell Isolation Kit (Stemcell), per manufacturer’s instructions. Bulk T cells were cultured in ImmunoCult-XF T Cell Expansion Medium (Stemcell). Immediately after isolation, T cells were either frozen or stimulated for 2 days with anti-human CD3/CD28 magnetic Dynabeads (ThermoFisher) at a bead to cell concentration of 1:1, along with cytokine 500U/mL IL-2 (Peprotech), 5 ng/mL IL-7 (Peprotech), and 5 ng/mL IL-15 (Peprotech). Following stimulation, cells were maintained at 50 U/mL IL-2 with the addition of fresh media every 2 days. Cells were always maintained at a density of 1 million cells per mL.

### microRNA eIF3 RIP-qPCR

EIF3B immunoprecipitation (IP) was performed using 1 × 10⁷ primary T cells per condition. When indicated, cells were activated with anti-human CD3/CD28 magnetic Dynabeads (ThermoFisher) for 5h prior to lysis. Unactivated cells were left untreated and processed directly for lysis. For activated samples, T cell-bead complexes were detached by scraping and gently pipetted to dissociate prior to lysis. All samples were then lysed on-bead under identical conditions. Fresh cell lysis buffer (50 mM HEPES-KOH pH 7.5, 150 mM KCl, 2 mM EDTA, 0.5 mM DTT, 0.5% NP-40, EDTA-free protease inhibitor, and 11 µL murine RNase inhibitor (NEB) per mL) was prepared, and 1 mL was added per 1 × 10⁷ cells. Pellets were resuspended by gentle pipetting, incubated on ice for 10 min, and lysed by passing through an 18-gauge needle five times before centrifugation at 13,000 × g for 15 min at 4 °C. Clarified supernatants were aliquoted for immunoprecipitation. For IP, 100 µL Protein G Dynabeads per 1 × 10⁷ cells (Invitrogen) were washed twice in cold lysis buffer and incubated with 5 µg anti-EIF3B antibody (Bethyl Laboratories) or control IgG (Cell Signaling Technology) for 45 min at room temperature with rotation. Antibody-coupled beads were washed twice and incubated with 1 mL of clarified lysate per 1 × 10⁷ cells for 2 h at 4 °C with rotation. Beads were magnetically separated, and the supernatant was collected as western blot (WB) flow-through, if needed. Beads were then washed three times in cold low-salt wash buffer (50 mM HEPES-KOH pH 7.5, 150 mM KCl, 0.5 mM DTT, 0.5% NP-40, EDTA-free protease inhibitor, and 11 µL murine RNase inhibitor per mL) with 5 min incubations at 4 °C. We tested three independent methods for RNA isolation from the eIF3 RIP experiments: Zymo total RNA (Direct zol RNA Miniprep R2051), Zymo small RNA enrichment (Quick RNA Microprep kit R1050), and miRVana small RNA enrichment (Thermofisher AM1561).

All three methods yielded RNA suitable for qPCR analysis. However, across isolation methods, the miRVana small enrichment protocol was the most robust, followed by Zymo small RNA enrichment. We used qPCR primers designed specifically to amplify a subset of miRNAs of interest (Thermofisher TaqMan Advanced miRNA Assays), and used these primers across all RNA isolations. For RNA isolation (1 × 10⁷ cell equivalents), beads were resuspended in 500 µL TRI Reagent or 300 µL miRVana lysis buffer, vortexed, magnetically separated, and the resulting supernatant was collected for RNA extraction using either the Zymo Directzol RNA/ small RNA enrichment kit or the mirVana miRNA isolation kit (Invitrogen), respectively as per manufacturer’s instructions. For mirVana samples, cells were treated with Turbo DNase kit (Ambion) according to manufacturers’ instructions. For WB analysis, beads were incubated at 70 °C for 10 min in 50 µL NuPAGE loading dye (Invitrogen), vortexed, magnetically separated, and the supernatant was collected for western blotting. miRNA qPCR was carried out with Taqman Advanced miRNA assays as per manufacturer’s instructions (ThermoFisher).

### Jurkat cell culture

Human Jurkat Clone E6-1 (ATCC TIB-152) was obtained from the Berkeley Cell Culture Facility. Cells were routinely tested for mycoplasma contamination at the UC Berkeley Cell Culture Facility. Cells were maintained in RPMI 1640 Medium (ATCC modification) with 10% FBS (VWR Seradigm Life Science) and 0.01% Penicillin-Streptomycin (10,000 U/mL) (ThermoFisher). The cells were maintained between 1 x 10^5^ cells mL^−1^ to 1 × 10^6^ cells mL^−1^. Where indicated, Jurkat cells were activated with plate-bound anti-CD3 and soluble anti-CD28 antibodies.

### Jurkat cell miRNA editing

Jurkat E6.1 cells were maintained in RPMI-1640 supplemented with 10% FBS and used for genome editing at low passage number. CRISPR-Cas9 ribonucleoprotein (RNP) complexes were assembled immediately prior to nucleofection. For each reaction, 1 µL Cas9 protein (40 µM) and 1 µL of sgRNA (100 µM; Synthego) were thawed rapidly, combined, and the volume was adjusted to 10 µL using supplemented SE Cell Line Nucleofector Solution (Lonza), which was also used for subsequent cell resuspension. Cas9 was added slowly with gentle circular mixing to promote complex formation, and RNPs were incubated at 37 °C for 15 min and kept at room temperature until use, followed by addition of 1 µL of 100 µM HDR donor template (IDT) when applicable. Jurkat cells were counted, and 0.5-1 × 10⁶ cells were used per nucleofection reaction. Cells were pelleted, washed once with PBS, and resuspended in supplemented SE Cell Line Nucleofector Solution (16.4 µL P3 + 3.6 µL Supplement per reaction). For each reaction, 20 µL of the cell suspension was combined with 10 µL of the RNP mixture, and 30 µL was transferred into a well of a 4D-Nucleofector 16-well shuttle plate while avoiding air bubbles. Nucleofection was performed using program CL-120, after which 80–100 µL of pre-warmed RPMI-1640 medium containing 10% FBS was added directly to each well. Cells were then transferred to a 24-well plate and cultured under standard conditions for recovery and subsequent analysis.

### Lentiviral production, transduction and rescue experiments in Jurkat cells

For lentiviral production, HEK293T cells were plated at a density of 80% in T-75 flasks the night before transfection. The cells were then transfected with the plasmid expressing the protein of interest (CD813A-System Biosciences), PsPAX2 and pCMV-VSV-G using the Lipofectamine 2000 reagent (ThermoFisher) following the manufacturer’s instructions. 48 and 72 hours after transfection, the viral supernatant was collected, filtered and concentrated using PEG-it Virus Precipitation Solution (System Biosciences) following the manufacturer’s instructions. The virus pellets were then resuspended in RPMI-1640 media supplemented with 25 mM HEPES at approximately 100× the original concentration, transferred to cryogenic vials, and stored in -80 °C.

For each lentivirus transduction reaction, 5 × 10⁵ cells were resuspended in a transduction mixture consisting of 400 μL complete RPMI medium, 100 μL MAX Enhancer (System Biosciences), 2 μL TransDux (System Biosciences), and 4 μL 1 M HEPES buffer (final HEPES concentration 8 mM). A total of 506 μL of the cell-enhancer-TransDux mixture was added to each well of a 24-well plate, followed by 50 μL of lentiviral particles encoding the indicated constructs (Control and the miR-17∼92 cluster). Plates were centrifuged at 1,500 × g for 1.5-2 h at 32 °C to facilitate spinoculation. After centrifugation, 400 μL complete medium was added to each well, cells were gently mixed by pipetting, transferred to microcentrifuge tubes, and centrifuged at 1,500 × g for 5 min. Pellets were resuspended in 400 μL fresh complete RPMI medium (without TransDux) and transferred to new wells of a 24-well plate. Cells were incubated at 37 °C in a 5% CO₂ incubator for downstream assays.

For rescue experiments, the miR-17∼92 cluster was cloned into CD813A lentiviral vectors (System Biosciences) under the control of a common EF1α promoter, with GFP expressed from a PGK promoter and an SV40 poly(A) signal. The corresponding empty vector served as the control construct.

### mRNA sequencing in Jurkat cells

RNA samples were extracted from unactivated or 5 h activated edited (KO-Clone1 or KO-Clone2) and unedited (WT) Jurkat clones, using the Direct-zol RNA Miniprep kit (Zymo Research) and submitted to Novogene for library preparation and deep sequencing. We performed bulk RNA-seq on NovaSeq PE150 paired-end libraries and first verified data integrity by MD5 checksums, followed by quality assessment with FastQC (v0.12.1). Adaptor and low-quality base trimming were carried out with cutadapt (Martin, 2011) using Illumina TruSeq adaptor sequences and a Phred quality cutoff of Q20, and trimming was repeated for all samples in a shell loop, with only trimmed FASTQ files retained for downstream analysis. Transcript quantification was done with kallisto (Bray et al., 2016) using an Ensembl GRCh38 cDNA reference (Homo_sapiens.GRCh38.cdna.all.fa.gz) to build the index, with 100 bootstrap samples per library to capture technical variance. Differential expression was performed in R with sleuth (Pimentel et al., 2017) at the transcript level, using a design ∼ genotype (WT vs KO-Clone1 or KO-Clone2) analyzed separately at 0 h and 5 h, with WT as the reference. Wald tests provided log_2_ fold changes (b) and FDR-adjusted q-values; transcripts were considered differentially expressed at qval < 0.05 and |log2FC| > 1 (or > 2 for stricter subsets). Transcript IDs (target_id) were mapped to gene symbols by parsing the Ensembl cDNA FASTA headers, and transcript-level results were collapsed to gene-level by keeping, per gene and direction, the isoform with the largest |log2FC|.

Overlapping DE genes between KO-Clone1 and KO-Clone2 were defined at each timepoint and stratified into up- and downregulated sets, which were used for Venn diagrams, T-cell marker panels (based on a curated list including IL2, IFNG, IL2RA, CD44 and other activation/checkpoint genes), and log2FC bar/volcano plots. Gene Ontology Biological Process enrichment on overlapping 5 h gene sets (downregulated, and combined up+down) was performed with clusterProfiler and org.Hs.eg.db using gene symbols, Benjamini-Hochberg correction, and an adjusted p-value (FDR) cutoff of 0.05; enriched terms were visualized as dot plots generated in ggplot2.

### Primary T cell miRNA editing

Freshly isolated or thawed primary human CD3⁺ T cells were activated with CD3/CD28 beads (ThermoFisher) for 2 days prior to genome editing. After activation, cells were pelleted and prepared for CRISPR-Cas9 nucleofection using pre-assembled ribonucleoprotein (RNP) complexes. For each reaction, 1 µL Cas9 protein (40 µM) and 1 µL of each sgRNA (100 µM; Synthego) were thawed rapidly, combined, and the volume was adjusted to 10 µL using supplemented Lonza P3 nucleofector solution (Lonza), which was also used later for resuspending cells. Cas9 was added slowly with gentle circular mixing to facilitate complex formation. RNPs were incubated at 37°C for 15 min and then kept at room temperature until use, followed by addition of 1 µL of 100 µM HDR donor template (IDT) when applicable. Activated cells were counted, and 0.5-1 × 10⁶ cells were used per nucleofection reaction; cells were washed with PBS and resuspended in P3 buffer supplemented according to the manufacturer’s instructions (16.4 µL P3 + 3.6 µL Supplement per reaction). For each reaction, 20 µL of the cell suspension was mixed with 10 µL of the RNP mixture, and 30 µL was transferred into each well of a 4D-Nucleofector 16-well shuttle plate, avoiding bubbles. Nucleofection was performed using program EH115, after which 80–100 µL of pre-warmed primary T cell medium supplemented with 50 U/mL IL-2 was added directly to each well, and cells were transferred to 12-well plates for recovery and subsequent culture.

### gDNA isolation-PCR

48 hours after nucleofection of Jurkat and primary T cells, genomic DNA was isolated using the Quick Extraction DNA Extraction Solution (Epicentre) according to the manufacturer’s guidelines with minor modifications. Briefly, washed cell pellets were resuspended in 100 μL Quick Extraction solution, incubated at room temperature for 5 min, mixed by gentle pipetting and a brief 15s vortex, and subsequently incubated at 65 °C for 15 min followed by 98 °C for 10 min to complete lysis. Genomic DNA concentration was measured using the Quick Extraction solution as the blank, and samples were stored at -20 °C until use. Flanking primers were used to PCR-amplify the targeted regions, including the deleted loci in knockout clones and the corresponding non-deleted regions in scrambled sgRNA controls (Supplementary Table 3). PCR products were purified and submitted to Plasmidsaurus for amplicon sequencing.

### Nanopore amplicon sequencing and deletion quantification

PCR amplicons spanning the edited *MIR17HG* locus were purified and submitted for Nanopore amplicon sequencing (Plasmidsaurus PCR Premium). Reads were aligned to the wild-type amplicon reference sequence using BWA-MEM and visualized in IGV. Deletion efficiency was quantified using a length-based approach by classifying reads <900 bp as deleted and reads >1200 bp as WT-like, excluding intermediate-length reads. Percent deletion was calculated as the fraction of deleted reads among informative reads.

### Flow cytometry and sorting

Flow cytometric analysis was conducted using an Attune NxT Acoustic Focusing Cytometer (ThermoFisher). For surface staining and cell sorting, cells (Jurkat and primary T cells) were pelleted and resuspended in 50 μL of FACS buffer (2% FBS in PBS) containing antibodies (CD25-APC and CD69-PE) at a 1:100 dilution (BioLegend) and incubated on ice in the dark for 30 minutes. Following staining, cells were washed twice in the FACS buffer, resuspended, and analyzed. To assess cell surface expression of CD25 and CD69, gating for CD25^+^CD69^+^ cells was established based on CD25 and CD69 levels in non-activated T cells. For Figure 5, cells were stained with Ghost Dye Violet 450 (Cytek Biosciences) and CD25-APC (BioLegend) in 50 µL staining buffer (1:100 antibody dilution in FACS buffer) per 1 × 10⁶ cells. At least 100,000 events were recorded, and cells were sorted using a Sony Biotechnology cell sorter. All flow cytometry data were processed using FlowJo software (BD Life Sciences).

### Mass spectrometry of eIF3 IP samples

EIF3B IP was performed using unactivated and 5h activated primary T cells as described above in the microRNA eIF3 RIP-qPCR methods section. For RNase I treated samples, RNase I (Ambion) was added at 5 U per 1 × 10^7^ cells prior to immunoprecipitation. For mass spectrometry, 25 µL of each EIF3B IP eluate was loaded onto a precast SDS-PAGE gel and run until proteins entered the resolving gel. Gels were washed, stained with GelCode Blue (Thermofisher Scientific) for 1 h, destained with Milli-Q water, and protein bands were excised and submitted for mass spectrometry. Raw MS1 peak area values (“Area”) were obtained for three independent donors, each analyzed as paired IgG control and EIF3B IP samples at 0 h and 5 h, with and without RNase treatment. Area values were merged by gene symbol across donors, log2-transformed as log2(Area + 1), and analyzed using limma (Hutchings et al., 2023; Ritchie et al., 2015) with the design ∼ IP + Donor. Missing values were set to zero prior to log2 transformation. IgG served as the reference, and donor was included as a blocking factor. Positive log_2_ fold changes reflect enrichment in eIF3B IP relative to IgG. This yielded donor-corrected log_2_ fold changes and Benjamini–Hochberg–adjusted p-values for EIF3B enrichment relative to IgG. GO Biological Process enrichment was performed on donor-corrected EIF3B IP datasets using clusterProfiler (Wu et al., 2021). For each 5 h condition (±RNase), eIF3 complex subunits (EIF3A-EIF3M) were excluded, and proteins with log_2_ fold change > 1 and adjusted p-value < 0.05 were used as the foreground set. All quantified proteins served as the background. GO terms were identified using a hypergeometric test with Benjamini-Hochberg correction and visualized using enrichplot.

## Data Availability Statement

Data underlying this article are publicly available. mRNAseq data are deposited under GSE328211. The code for data processing is available upon request.

## Funding

This work was supported by the National Institutes of Health (NIH) grants R01-GM065050 and R35-GM148352 (to J.H.D.C).

## Competing interests

Authors declare no competing interests.

## Contributions

P.M. and J.H.D.C. designed the study. P.M. performed all molecular biology experiments, data analysis for mRNAseq and IP-MS, with preliminary Ago2 data from D.D.S. that guided IP-MS experiments in primary T cells. J.S. and C.H. assisted P.M. with RIP-qPCR experiments. C.Y.C. assisted P.M. with the rescue experiment. Y.K. performed Ago2 western blot analyses. A.H.B. assisted J.H.D.C. with reanalysis of PAR-CLIP data. Previously published PAR-CLIP data generated by D.D.S. were reprocessed in this study. P.M. and J.H.D.C. wrote the manuscript, with input from J.S., C.H., and Y.K. All authors approved the final version of the manuscript.

## Acknowledgements

We thank the members of the J.H.D.C. laboratory for the helpful discussion. Figures 2A, 3A, 5A, 6A and 7 were created in https://BioRender.com.

## Figure Supplements

**Figure 1–figure supplement 1.**
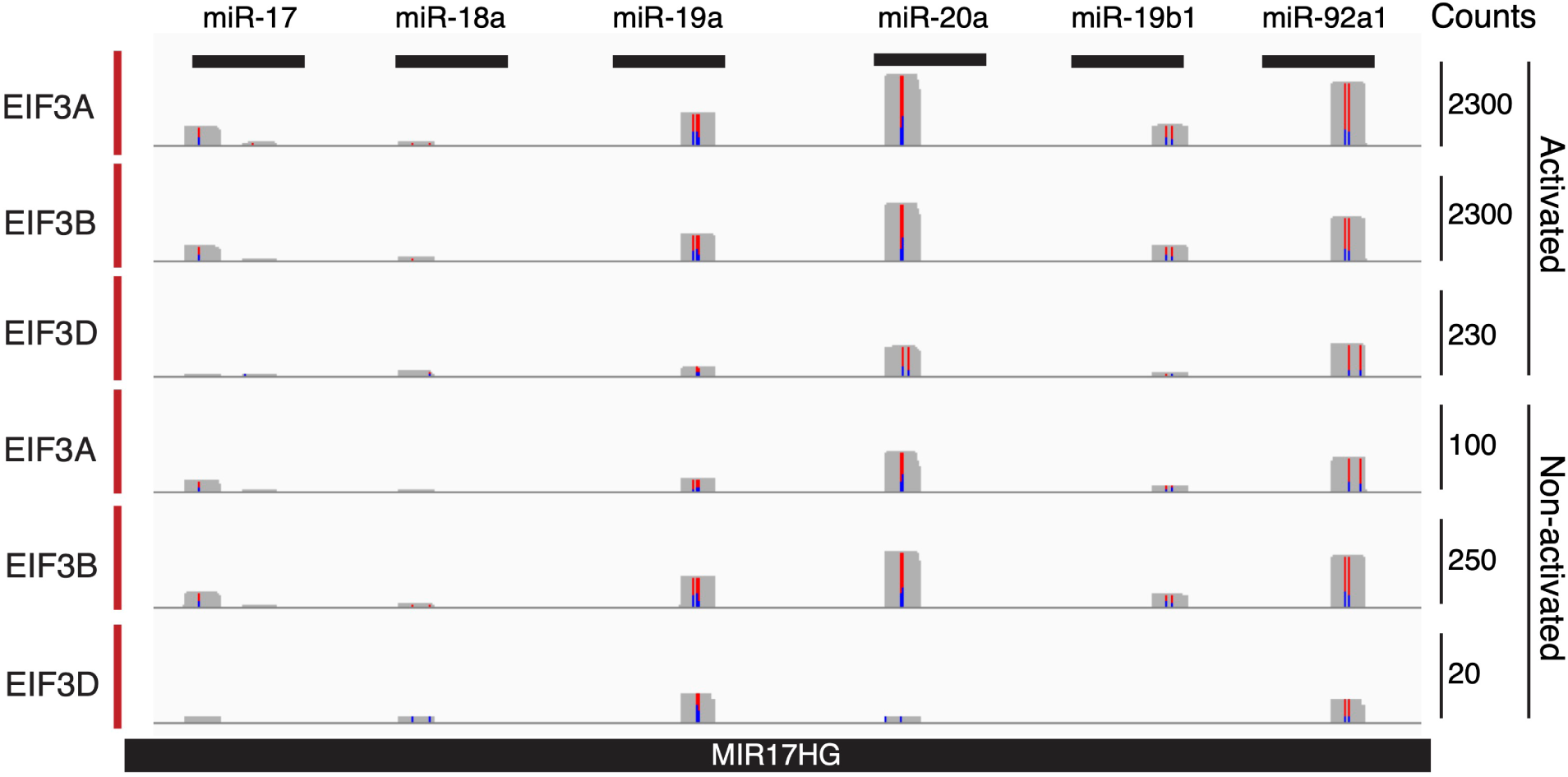
PAR-CLIP analysis of eIF3 subunit crosslinking to the miR-17∼92 cassette in Jurkat cells. PAR-CLIP signal tracks showing crosslinking of EIF3A, EIF3B, and EIF3D across individual mature miRNAs within the miR-17∼92 cassette in activated (5 hour) and unactivated Jurkat cells (replicate 2).

**Figure 1–figure supplement 2.**
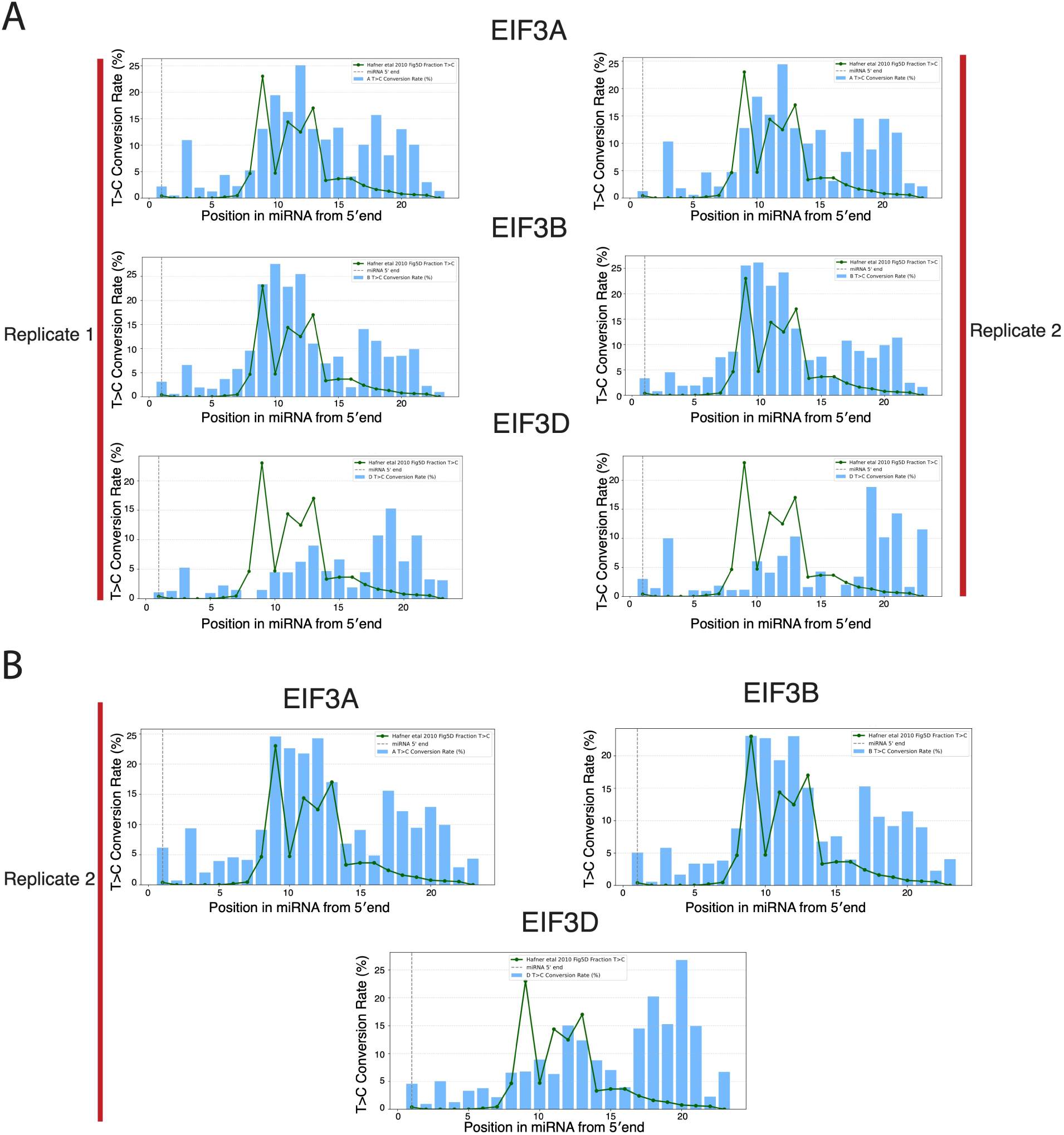
miRNA binding patterns with eIF3 subunits in Jurkat cells. (A) T>C conversion profiles from PAR-CLIP data plotted relative to the miRNA 5′ end for EIF3A, EIF3B, and EIF3D in unactivated Jurkat cells, shown for two independent replicates. (B) T>C conversion profiles from PAR-CLIP data plotted relative to the miRNA 5′ end for EIF3A, EIF3B, and EIF3D in activated Jurkat cells (replicate 2). Blue bars represent the T>C conversion rate (%) at each nucleotide position along the miRNA, and the green line represents the published AGO2 T>C conversion profile from Hafner *et al*. (2010) and the grey dotted line represents the miRNA 5′ end.

**Figure 1–figure supplement 3.**
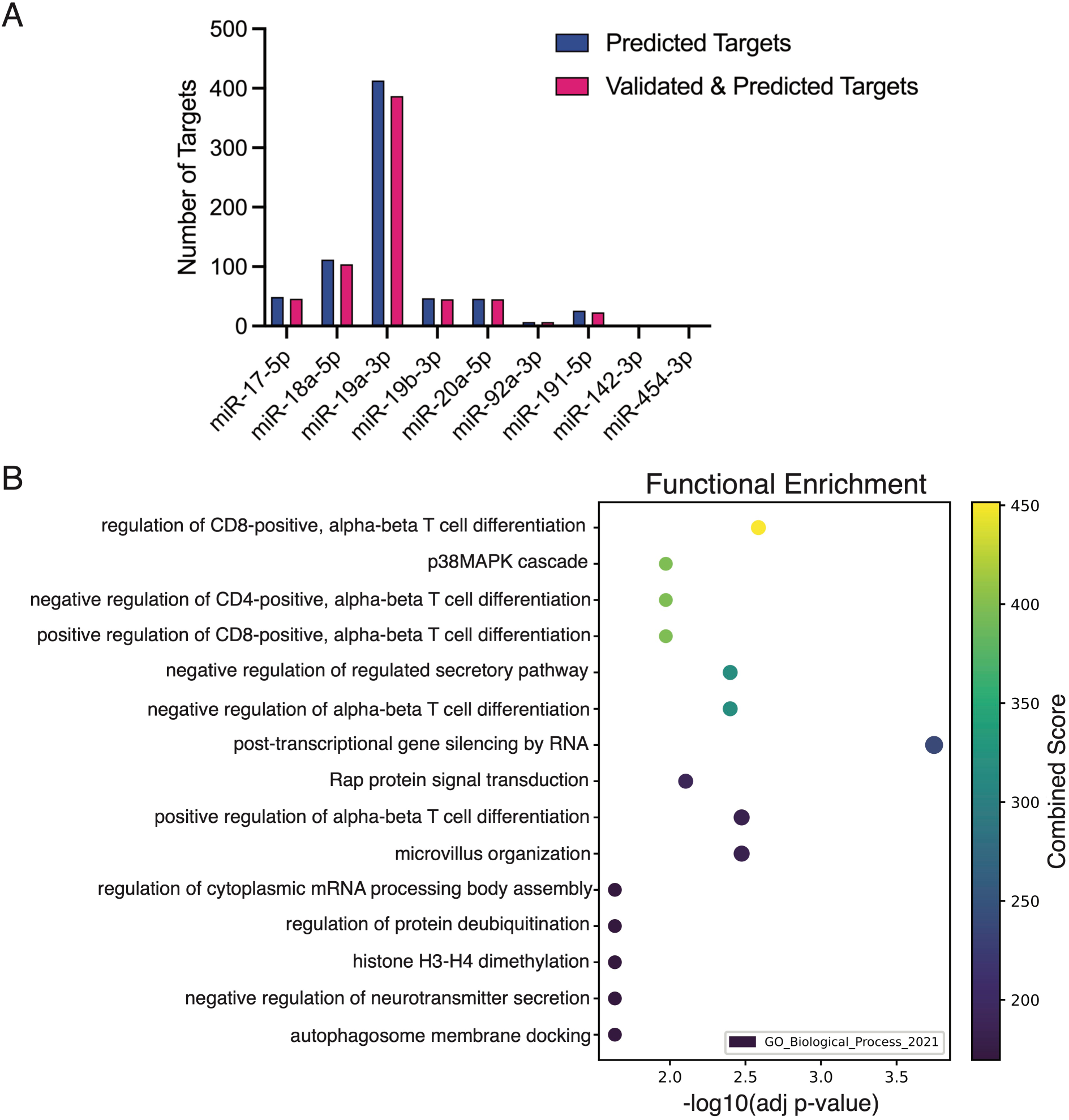
Overlap of miR-17∼92 predicted and validated targets with eIF3-mRNA crosslinking data. (A) Number of “predicted” and “predicted and validated” miR-17∼92 target mRNAs that overlap with eIF3-mRNA Jurkat PAR-CLIP data, shown for individual miRNAs within the miR-17∼92 cluster. (B) Gene Ontology analysis of overlapping target genes identified in panel A.

**Figure 2–figure supplement 1.**
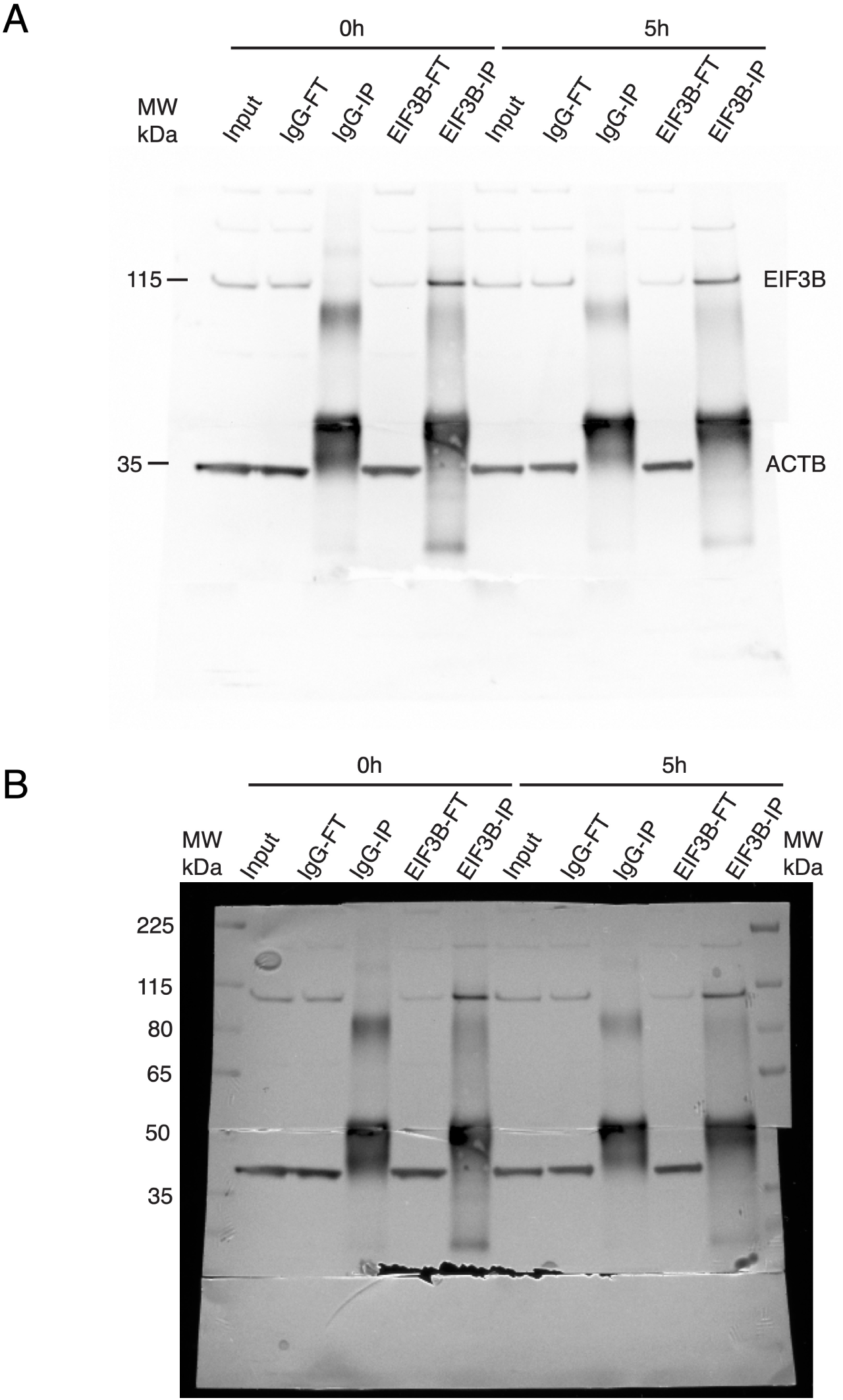
Uncropped western blot images corresponding to Figure 2. Unprocessed western blot images for EIF3B immunoprecipitation experiments shown in Figure 2B. Top and bottom blots are two images of different exposure times of the same blot.

**Figure 2–figure supplement 2.**
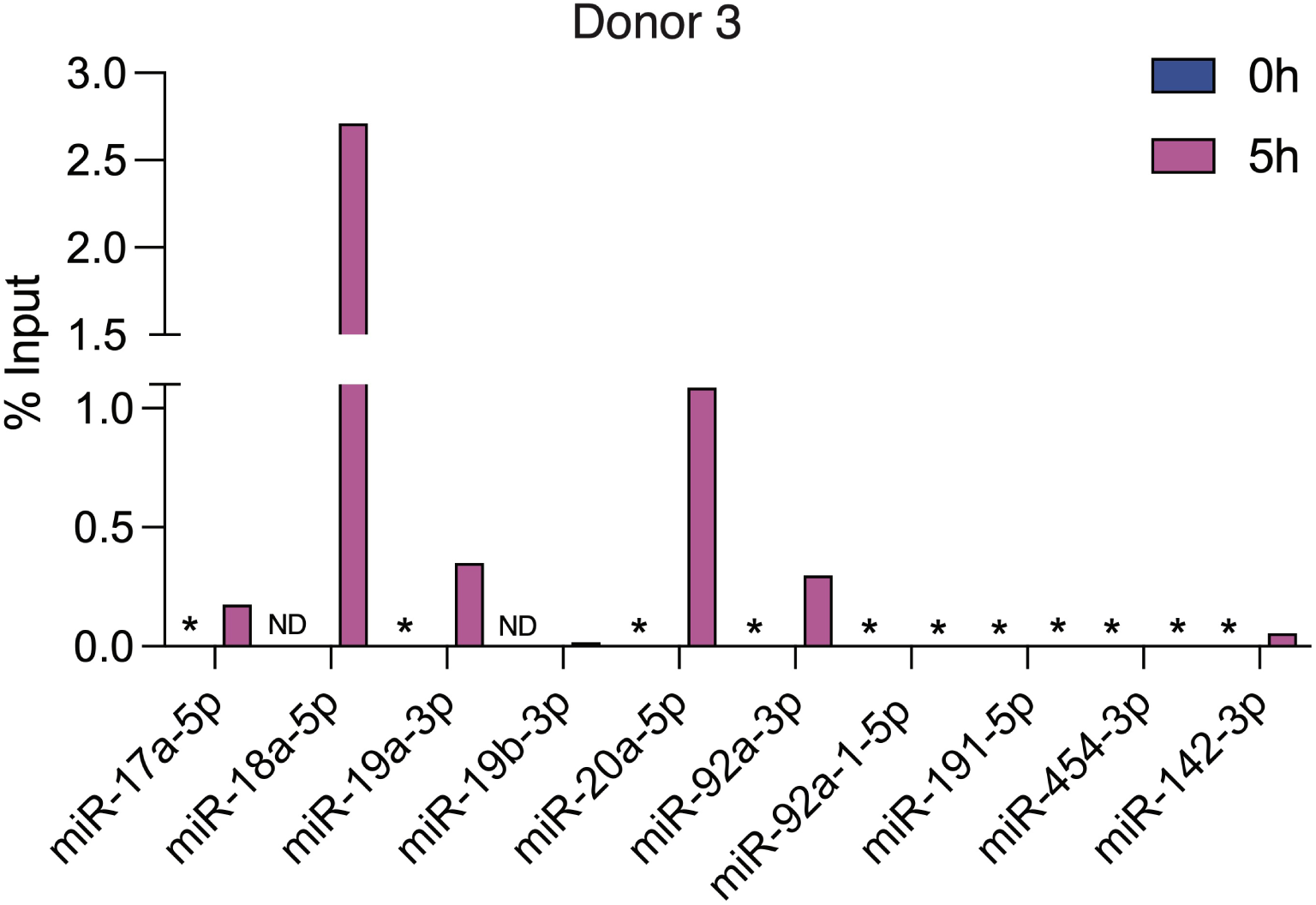
RIP-qPCR analysis of EIF3B-associated miRNAs in primary T cells. RIP-qPCR analysis of miRNAs from the miR-17∼92 cassette enriched in EIF3B immunoprecipitation from primary T cells activated for 5 hour (Donor 3), plotted as % Input (% Input EIF3B minus % Input IgG). Asterisk denotes samples where IgG or EIF3B IP Cq values were at or above the detection threshold; ND, not detected in either input or IP samples.

**Figure 3–figure supplement 1.**
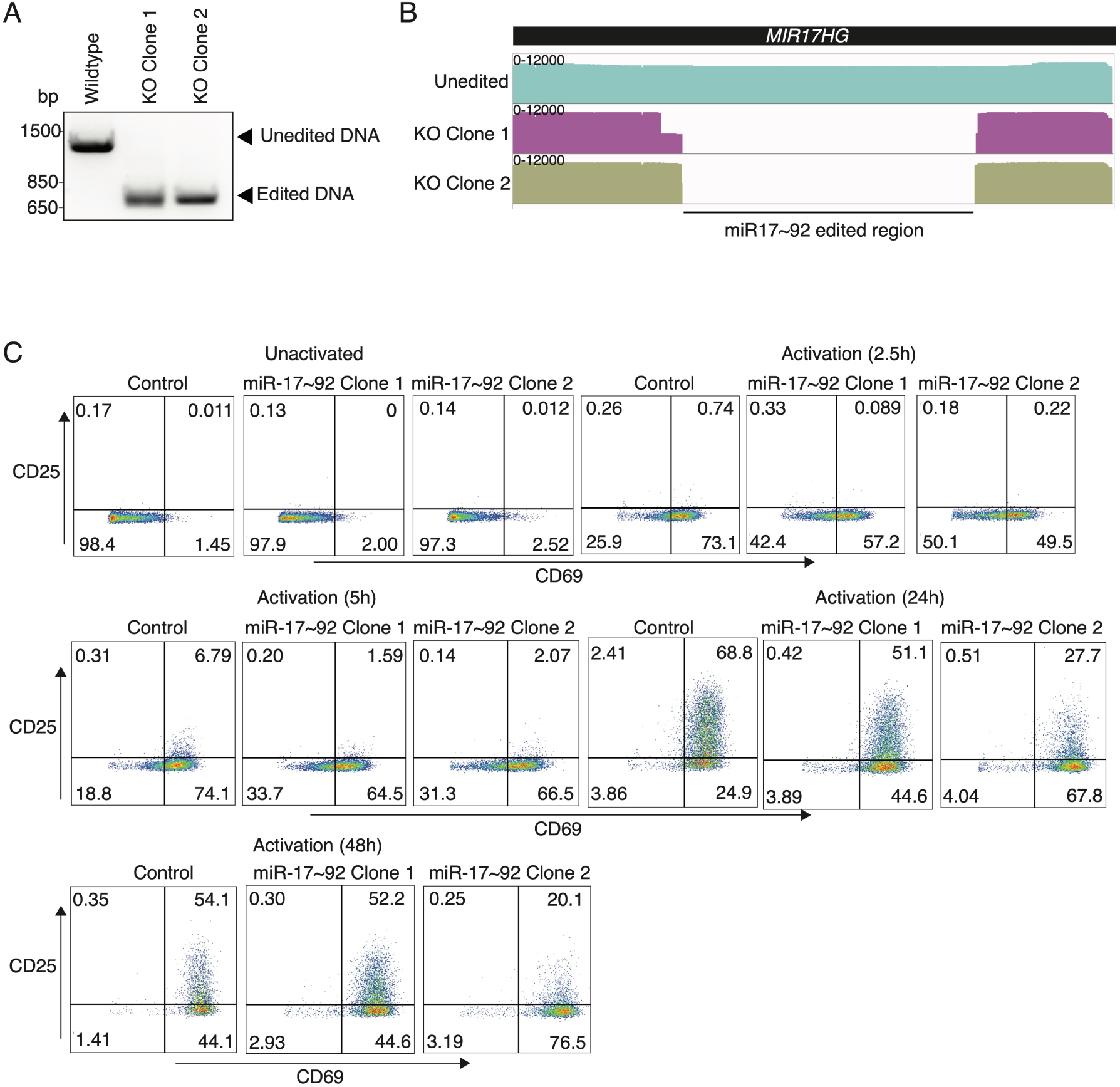
Additional characterization of miR-17∼92 knockout Jurkat cells. (A) Genotyping PCR confirming deletion of the miR-17∼92 locus in knockout clones. (B) IGV visualization of long read amplicon sequencing data confirming edits at the miR-17∼92 locus in knockout clones. (C) Flow cytometry plots showing CD69 and CD25 expression in control and miR-17∼92 knockout clones under unactivated conditions and following T cell activation at the indicated time points.

**Figure 3–figure supplement 2.**
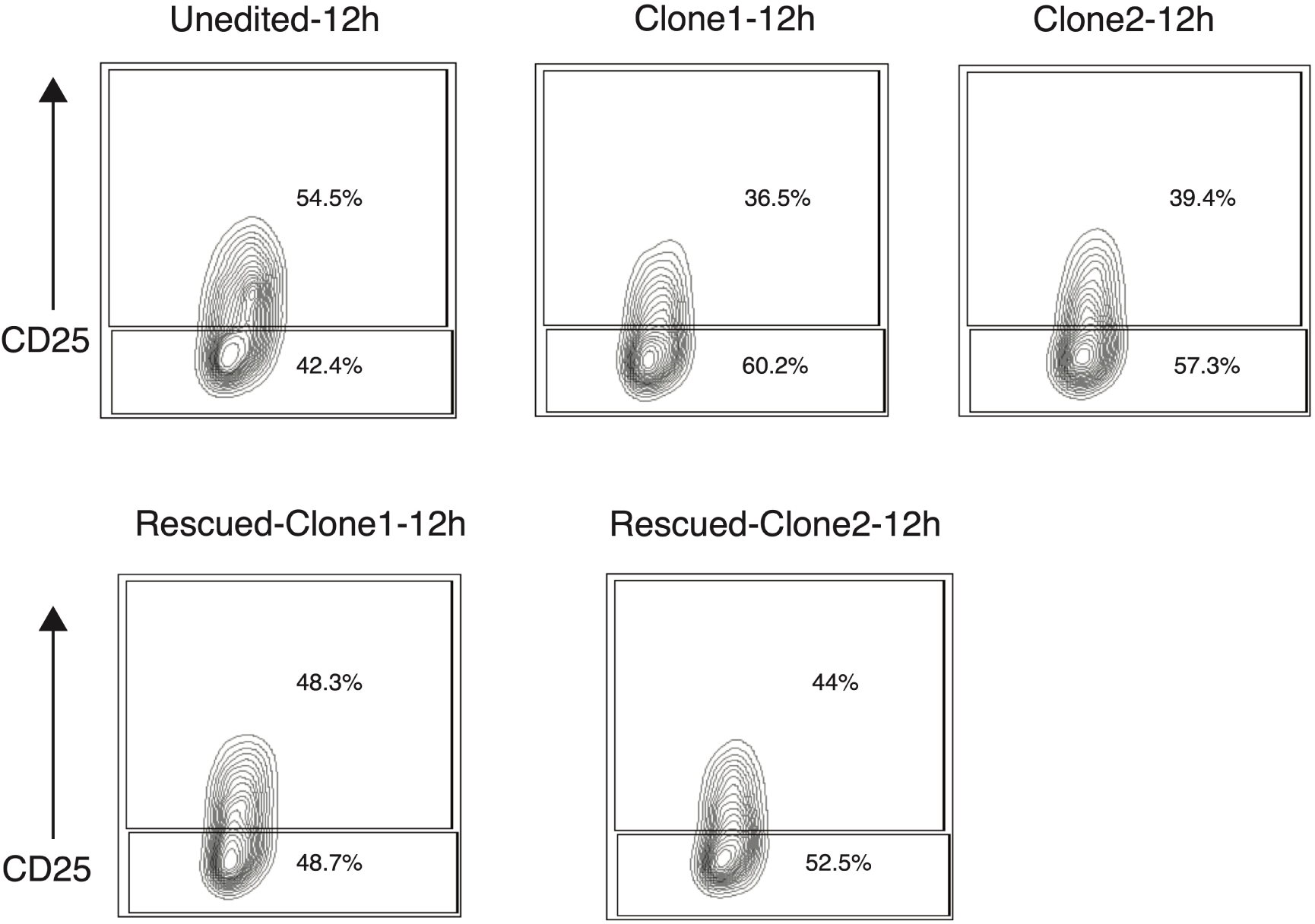
Rescue of miR-17∼92 knockout phenotype in Jurkat cells. Flow cytometry plots showing CD25 expression in unedited Jurkat cells, miR-17∼92 knockout clones, and rescued clones following 12 hours of activation. Percentages indicate the fraction of cells within each gate.

**Figure 4–figure supplement 1.**
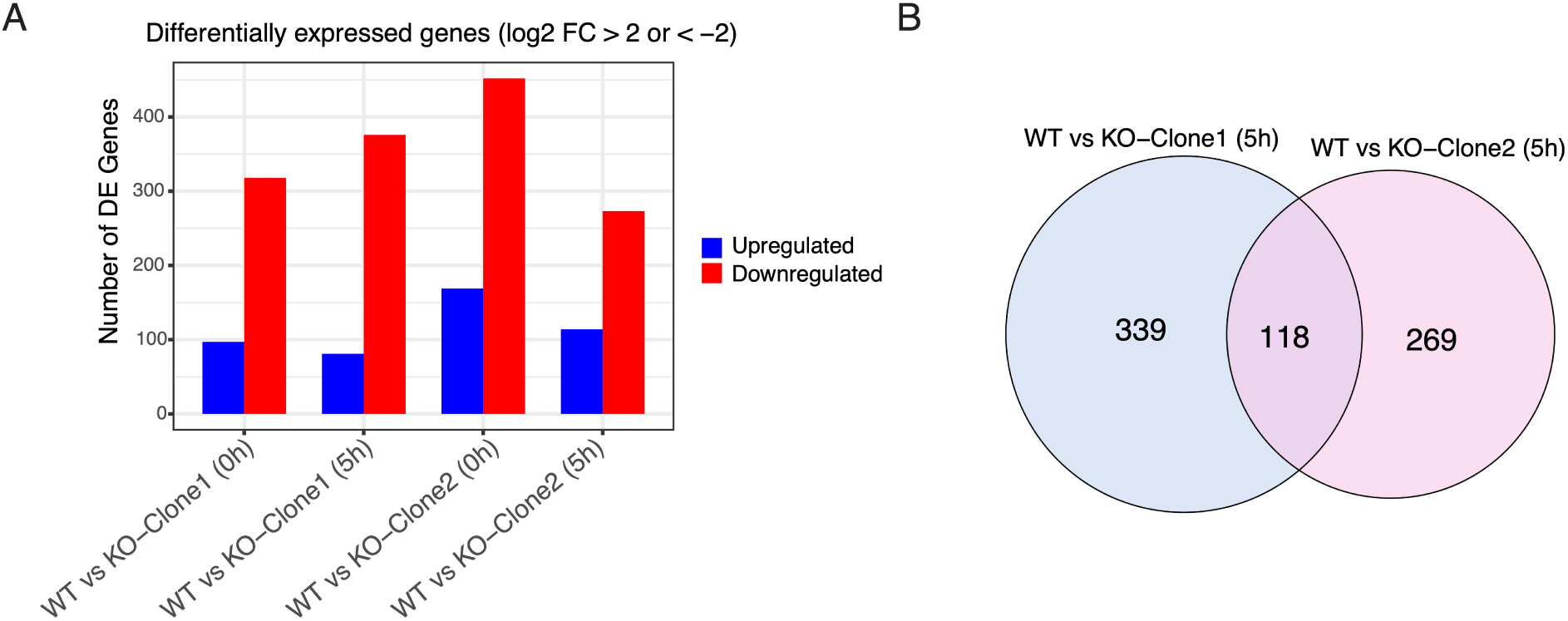
Differential gene expression in miR-17∼92 knockout clones. (A) Number of differentially expressed genes (|log_2_ fold change| > 2) in miR-17∼92 knockout clones relative to wild-type Jurkat cells under unactivated and 5 hour activation conditions, separated into upregulated and downregulated genes. (B) Overlap of differentially expressed genes (|log_2_ fold change| > 2) between miR-17∼92 knockout clone 1 and clone 2 at 5 hours of activation.

**Figure 5–figure supplement 1.**
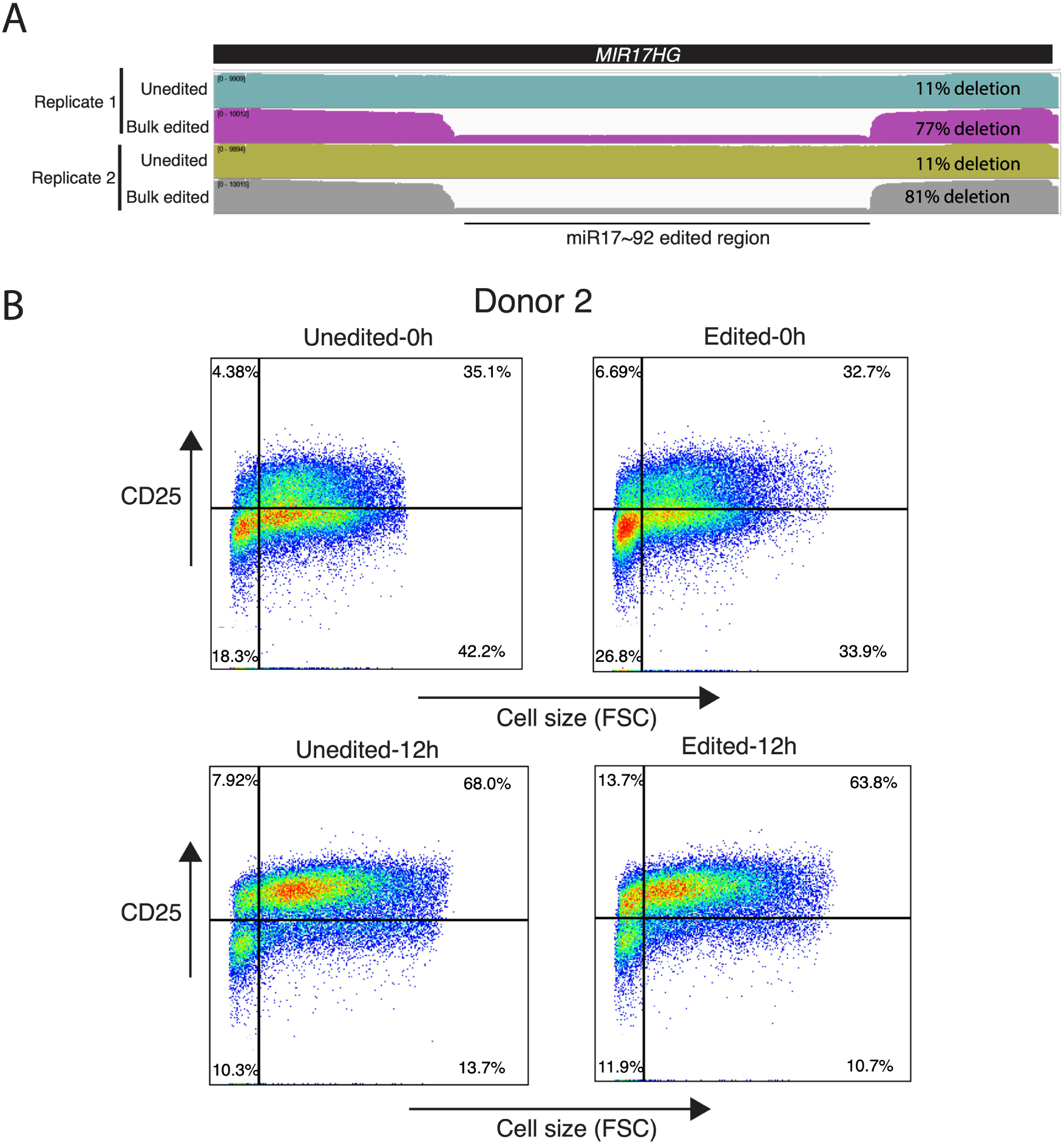
Validation of miR-17∼92 editing efficiency and phenotypic assessment in bulk-edited primary T cells. (A) IGV snapshots of long read amplicon sequencing data showing the miR-17∼92 edited region in unedited and bulk-edited primary T cells across two independent replicates, with indicated deletion efficiencies. Length-based quantification indicates deletion efficiencies of 77% and 81% in edited replicates, while control samples show background deletion signals of approximately 11%, likely reflecting alignment or PCR artifacts common in amplicon sequencing. (B) Flow cytometry analysis of CD25 expression and cell size in unedited and bulk-edited live primary T cells (Donor 2) under unactivated and 12 hour activation conditions.

**Figure 6–figure supplement 1.**
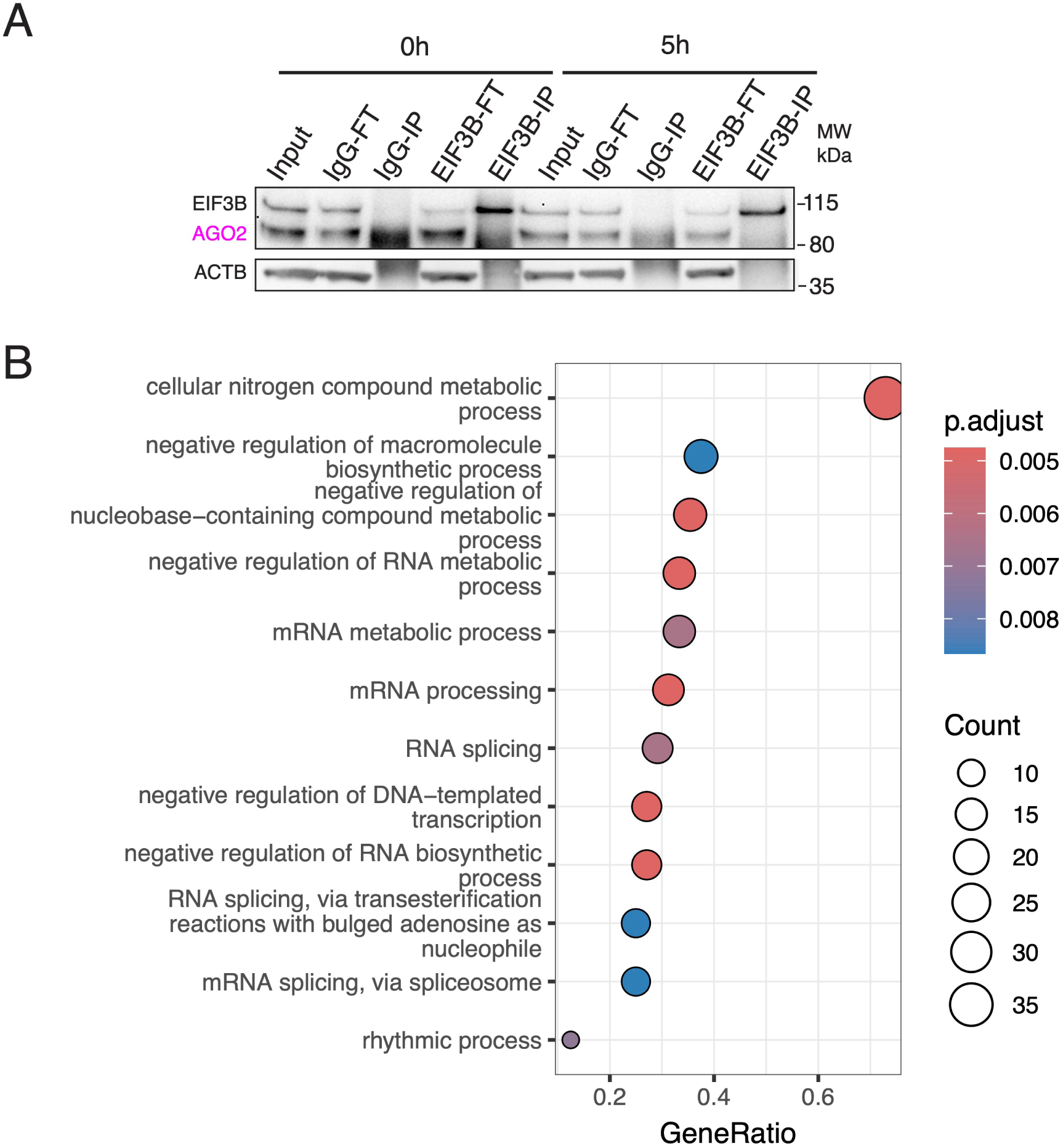
Lack of AGO2 detection in EIF3B immunoprecipitation from primary T cells. (A) Western blot analysis of AGO2 in EIF3B immunoprecipitation from primary T cells under unactivated and 5 hour activation conditions. Input, flowthrough (FT), and immunoprecipitated (IP) fractions are shown, with ACTB as a loading control. (B) Gene Ontology biological process (GO BP) enrichment analysis of EIF3B-associated proteins identified at 5 hour activation without RNase treatment.

## Notes

### Competing Interest Statement

The authors have declared no competing interest.

